# Predicting missing links in global host-parasite networks

**DOI:** 10.1101/2020.02.25.965046

**Authors:** Maxwell J. Farrell, Mohamad Elmasri, David Stephens, T. Jonathan Davies

**Affiliations:** Department of Biology, McGill University; Ecology & Evolutionary Biology Department, University of Toronto; Center for the Ecology of Infectious Diseases, University of Georgia; Department of Mathematics & Statistics, McGill University; Botany, Forest & Conservation Sciences, University of British Columbia; African Centre for DNA Barcoding, University of Johannesburg

## Abstract

1. Parasites that infect multiple species cause major health burdens globally, but for many, the full suite of susceptible hosts is unknown. Predicting undocumented host-parasite associations will help expand knowledge of parasite host specificities, promote the development of theory in disease ecology and evolution, and support surveillance of multi-host infectious diseases. Analysis of global species interaction networks allows for leveraging of information across taxa, but link prediction at this scale is often limited by extreme network sparsity, and lack of comparable trait data across species.
2. Here we use recently developed methods to predict missing links in global mammal-parasite networks using readily available data: network properties and evolutionary relationships among hosts. We demonstrate how these link predictions can efficiently guide the collection of species interaction data and increase the completeness of global species interaction networks.
3. We amalgamate a global mammal host-parasite interaction network (*>*29,000 interactions) and apply a hierarchical Bayesian approach for link prediction that leverages information on network structure and scaled phylogenetic distances among hosts. We use these predictions to guide targeted literature searches of the most likely yet undocumented interactions, and identify empirical evidence supporting many of the top “missing” links.
4. We find that link prediction in global host-parasite networks can accurately predict parasites of humans, domesticated animals, and endangered wildlife, representing a combination of published interactions missing from existing global databases, and potential but currently undocumented associations.
5. Our study provides further insight into the use of phylogenies for predicting host-parasite interactions, and highlights the utility of iterated prediction and targeted search to efficiently guide the collection of host-parasite interaction. These data are critical for understanding the evolution of host specificity, and may be used to support disease surveillance through a process of predicting missing links, and targeting research towards the most likely undocumented interactions.

## Introduction

Most disease-causing organisms of humans and domesticated animals can infect multiple host species (Cleaveland *et al*., 2001; Taylor *et al*., 2001). This has ramifications for biodiversity conservation (Smith *et al*., 2009) and human health via direct infection, food insecurity, and diminished livelihoods (Grace *et al*., 2012). Despite the severe burdens multi-host parasites can impose, we do not know the full range of host species for the majority of infectious organisms (Dallas *et al*., 2017). Currently, parasite host ranges are best described by host-parasite interaction databases that are largely compiled from primary academic literature. These databases gain strength by collating data across a large diversity of hosts and parasites, and have been used to identify macroecological patterns of infectious diseases (Stephens *et al*., 2016, 2017), define life histories influencing host specificity (Park *et al*., 2018; Park, 2019) and predict the potential for zoonotic spillover (Olival *et al*., 2017). However, global databases are known to be incomplete, with some estimated to be missing up to *∼*40% of host-parasite interactions among the species sampled (Dallas *et al*., 2017). While even incomplete data can offer important insights into the structure of host-parasite interaction networks, our ability to make accurate predictions of decreases when data are missing.

Filling in knowledge gaps, and building more comprehensive global databases of hostparasite interactions enhances our insights into the ecological and evolutionary forces shaping parasite biodiversity. These includes the role of network structure for transmission (Gomez *et al*., 2013; Pilosof *et al*., 2015), the nature of highly implausible host-parasite interactions (Morales-Castilla *et al*., 2015), the drivers of parasite richness (Nunn *et al*., 2003; Huang *et al*., 2015; Ezenwa *et al*., 2006; Kamiya *et al*., 2014) and parasite sharing across hosts (Davies & Pedersen, 2008; Huang *et al*., 2014; Braga *et al*., 2015; Luis *et al*., 2015; Albery *et al*., 2020), the association between host specificity and virulence (Shwab *et al*., 2018; Farrell & Davies, 2019), and global estimates of parasite diversity (Carlson *et al*., 2019). While determining the outcome of a given interaction ultimately requires moving beyond the binary associations provided by current databases, identifying documented and likely host-parasite associations is a critical first step.

Recent efforts have used species traits to predict reservoirs of zoonotic diseases (Han *et al*., 2015) and identify wildlife hosts of globally important viruses (Han *et al*., 2016; Pandit *et al*., 2018). Studies of global host-virus interactions have also been used to predict the structure of viral sharing networks (Albery *et al*., 2020; Wardeh *et al*., 2020). These approaches can identify host profiles for particular parasite groups, or host species that promote parasite sharing across hosts, but they do not predict interactions between individual host and parasite species. An alternative framework is to treat hosts and parasites as two separate interacting classes that can be represented in a bipartite network. Bipartite network models can predict new links based solely on the structure of the observed network, or incorporate node-level covariates, such as species traits (Dallas *et al*., 2017).

Algorithms such as recommender systems can identify probable links in a variety of large networks (Ricci *et al*., 2011). These models attempt to capture two key properties of real-world networks: the scale-free behaviour of interactions (shown by the degree distributions in Fig. SM 1) and local clustering of interactions (Watts & Strogatz, 1998). However, they also tend to predict that nodes with many documented interactions are more likely to associate with other highly connected nodes in the network (a behaviour described as “the rich get richer”). In the context of host-parasite interactions, the number of interactions per species will vary due to ecological or evolutionary processes, but may also be influenced by research effort. This is commonly seen in studies of parasite species richness in which sampling effort explains a large portion of the variation across hosts (Nunn *et al*., 2003; Ezenwa *et al*., 2006; Lindenfors *et al*., 2007; Kamiya *et al*., 2014; Olival *et al*., 2017). Models that directly or indirectly use the number of documented interactions per species to make predictions cannot easily adjust for these sampling biases. Thus, while these affinity based models may be highly tractable for large networks, they are also sensitive to uneven sampling across nodes, and may re-enforce observation biases.

Predictions from trait-informed network models are less sensitive to missingness, but require data on ecological and functional traits of interacting species (Dallas *et al*., 2017; Gravel *et al*., 2013; Bartomeus *et al*., 2016). Thus, while these approaches work well for smaller networks (Dallas *et al*., 2017), they can scale poorly to global-scale ecological datasets in which comparable traits are unavailable for all species (Morales-Castilla *et al*., 2015). In the absence of comprehensive trait data, we can incorporate evolutionary relationships among hosts to add biological realism to network-based predictions. Phylogenetic trees represent species’ evolutionary histories, and branch lengths of these trees provide a measure of expected similarities among species (Wiens *et al*., 2010). Hosts may be associated with a parasite through inheritance from a common ancestor, or as a result of host shifts (Page, 1993). In both processes we expect closely related species will host similar parasite assemblages (Davies & Pedersen, 2008).

Here we apply a link prediction model for bipartite ecological networks that combines properties of affinity based models with information on host phylogeny (Elmasri *et al*., 2020), to a massive global-scale host-parasite network for mammals. Incorporating host phylogeny into “rich-get-richer” (i.e. scale-free) interaction models adds biological structure, allowing for more realistic predictions with otherwise limited covariate data. This approach is particularly well-suited to link prediction in large global host-parasite networks as it does not require trait data, and allows accurate predictions of missing links in extremely sparse networks using only the structure of the observed host-parasite associations, and scaled evolutionary relationships among hosts. We demonstrate how model predictions can efficiently guide efforts to fill-in missing links, show their geographical distributions, and update existing global networks using historical and recently published interactions.

## Materials and Methods

### Data

To generate a network of documented host-parasite interactions for mammals, we amalgamated four major global host-parasite databases (Gibson *et al*., 2005; Wardeh *et al*., 2015; Stephens *et al*., 2017; Olival *et al*., 2017). These are derived from primary literature, genetic sequence databases, and natural history collections, and report host and parasite names as Latin binomials. The Global Mammal Parasite Database 2.0 (GMPD) (Stephens *et al*., 2016) contains records of disease causing organisms (viruses, bacteria, protozoa, helminths, arthropods, and fungi) in wild ungulates (artiodactyls and perrisodactyls), carnivores, and primates drawn from over 2700 literature sources published through 2010 for ungulates and carnivores, and 2015 for primates. The static version of the Enhanced Infectious Disease Database (EID2) (Wardeh *et al*., 2015) contains 22,515 host-pathogen interactions from multiple kingdoms based on evidence published between 1950 - 2012 extracted from the NCBI Taxonomy database, NCBI Nucleotide database, and PubMed citation and index. The Host-Parasite Database of the Natural History Museum, London (Gibson *et al*., 2005) contains over a quarter of a million host-parasite records for helminth parasites extracted from 28,000 references published after 1922, and is digitally accessible via the R package *helminthR* (Dallas, 2016). Finally, Olival *et al*. (2017) compiled a database of 2,805 mammal-virus associations for every recognized virus found in mammals, representing 586 unique viral species and hosts from 15 mammalian orders. These source databases were then combined into a single database, harmonising taxonomy of hosts and paratsites (full details in SM 1.1). We treat interactions as binary (0/1) for a given host-parasite pair as the sources do not explicitly indicate the role a host species plays in parasite transmission, but instead approximate host exposure and susceptibility to infection.

The resulting network includes 29,112 documented associations among 1835 host and 9149 parasite species (Fig. 1). To our knowledge this constitutes the largest mammal host-parasite interaction network currently available, and includes parasites from diverse groups including viruses, bacteria, protozoa, helminths, arthropods, and fungi, in wild, domestic, and human hosts. The resulting matrix is quite sparse, with *∼*0.17% of the *∼*16.8 million possible links having documented interactions. Humans are documented to associate with 2,064 parasites (47% of which associate with another mammal in the database), and comprise 7% of all interactions. Parasite species are largely represented by helminths (63.9%), followed by bacteria (13.1%) and viruses (7.89%). The degree distribution (number of documented interactions per species) varies considerably and is shown to be linear on the log scale for both hosts and parasites (Fig. SM 1).

**Figure 1:**
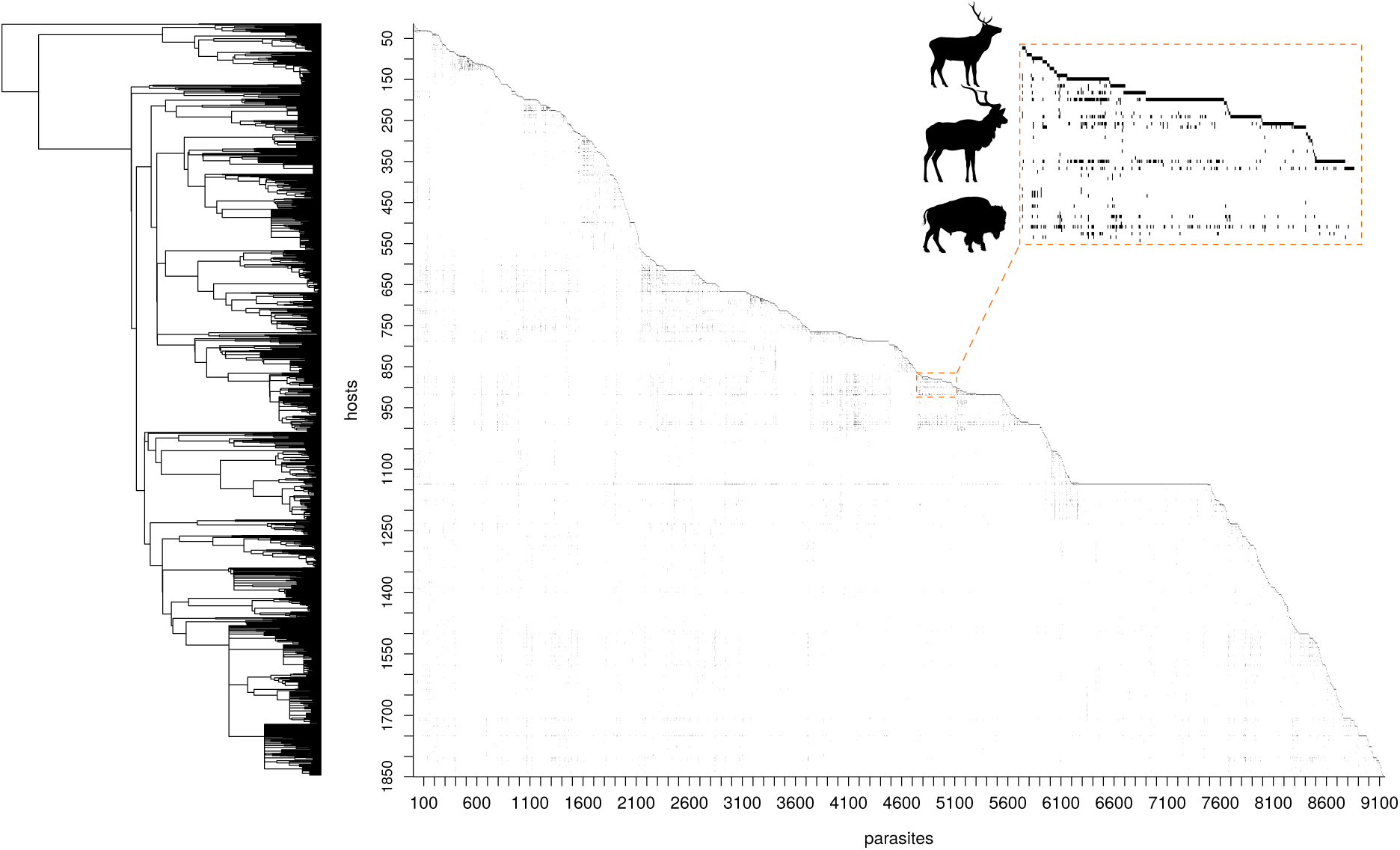
Phylogeny of host species (pruned from the Fritz et al. (Fritz *et al*., 2009) dated supertree for mammals), and host-parasite association matrix showing hosts ordered according to the phylogeny. Axes represent hosts (rows) and parasites (columns). The orange rectangles display an expanded subset of the matrix.

### Statistical analyses

We apply the bipartite link prediction model of Elmasri *et al*. (2020) to the amalgamated dataset. The model has three variants: the “affinity” model which generates predictions based only on the number of observed interactions for each host and parasite, the “phylogeny” model which is informed only by host evolutionary relationships (here taken from (Fritz *et al*., 2009)), and the “combined” model which layers both components (termed “full” model in Elmasri et al. (Elmasri *et al*., 2020)). The affinity model is fit by preferential attachment whereby hosts and parasites that have many interacting species in the network are assigned higher probabilities of forming novel interactions. The phylogeny model uses the similarity of host species based on evolutionary distances to assign higher probability to parasites interacting with hosts closely related to their documented host species, and lower probability of interacting with hosts that are distantly related. To account for uncertainty in the phylogeny, and allow the model to place more or less emphasis on recent versus deeper evolutionary relationships, we fit a tree scaling parameter (*η*) based on an accelerating-decelerating model of evolution (Harmon *et al*., 2010). This transformation allows for changes in the relative evolutionary distances among hosts and was shown to have good statistical properties for link prediction in a subset of the GMPD (El-masri *et al*., 2020). We apply these three models to the full dataset. The tree scaling parameter is applied across the whole phylogeny, but since the importance of recent versus deep evolutionary relationships among hosts is likely to vary across parasite types (Park *et al*., 2018), we additionally run the models on the dataset subset by parasite taxonomy (arthropods, bacteria, fungi, helminths, protozoa, and viruses). For all models we used 10-fold cross-validation to prevent over-fitting during parameter estimation, and to assess model performance when predicting links internal to the dataset (see SM 1.2 for details).

### Targeted literature searches

We identified the top 10 most likely links in each model-data subset combination which were not documented in the original data. These were used to guide searches of primary and grey literature for evidence of associations. Searches were conducted in Google Scholar by using both the host and parasite Latin binomials in quotes and separated by the AND boolean operator (e.g. “Gazella leptoceros” AND “Nematodirus spathiger”). If this returned no hits, the same search was conducted using the standard Google search engine in order to identify grey literature sources. For models run on the full dataset, we also investigate the top ten links for domesticated mammals (*Bison bison*, *Bos sp.*, *Bubalus bubalis*, *Camelus sp.*, *Capra hircus*, *Canis lupus*, *Cavia porcellus*, *Equus asinus*, *Equus caballus*, *Felis catus*, *Felis silvestris*, *Lama glama*, *Mus musculus*, *Oryctolagus cuniculus*, *Ovis aries*, *Rangifer tarandus*, *Rattus norvegicus*, *Rattus rattus*, *Sus scrofa*, *Vicugna vicugna*), and wild host species separately. For domesticated animals and humans, if the Latin binomials returned no hits, the search strategy was repeated using host common names (e.g. “human”, “pig”, “horse”).

We considered any physical, genetic, or serological identification of a parasite infecting a given host species as evidence of an association. Exceptions included 1) situations where parasite identification was stated as uncertain due to known serological cross-reactivity, 2) absence of clear genetic similarity to known reference sequences, or 3) unconfirmed visual diagnosis made from afar.

Missing links were classified as unlikely when involving ecological mismatch between host and parasite, such as trophically transmitted parasites infecting the wrong trophic level or unlikely ingestion pathway, and parasites typically classified as non-pathogenic or commensal. As our goal is to predict potential susceptibility, lack of known current geographic overlap among wide ranging hosts and parasites was not considered sufficient to classify a link as unlikely. In total, we generated 27 sets of “top ten” highly likely undocumented links to target, because of overlap in the top links among models run across data subsets, this reduced to 177 unique links investigated for published evidence of infection.

To contrast spatial predictions on the geographic distribution of missing links between the affinity and phylogeny models, we generated hotspot maps of undocumented host-parasite interactions. Maps were generated by summing the probabilities of undocumented host-parasite interactions per host species, summing across host species per cell (0.5 degree resolution) using IUCN distributions terrestrial mammals (IUCN, 2019). To adjust for the uneven species richness of hosts, predictions per cell were standardized to a scale of 0 - 1.

## Results

When assessing models using 10-fold cross-validation, all models performed well predicting links internal to the data. Area under the receiver operating characteristic curve (AUC) values ranged from 0.84 - 0.98 (maximum AUC of 1 signifies perfect predictive accuracy), and between 72.54% and 98.00% of the held-out documented interactions were successfully recovered (Fig. 2, Table SM 1; see Fig. SM 2 for posterior interaction matrices for the full dataset). Model performance tended to decrease with the size of the interaction matrix (Fig. 2A&C).

**Figure 2:**
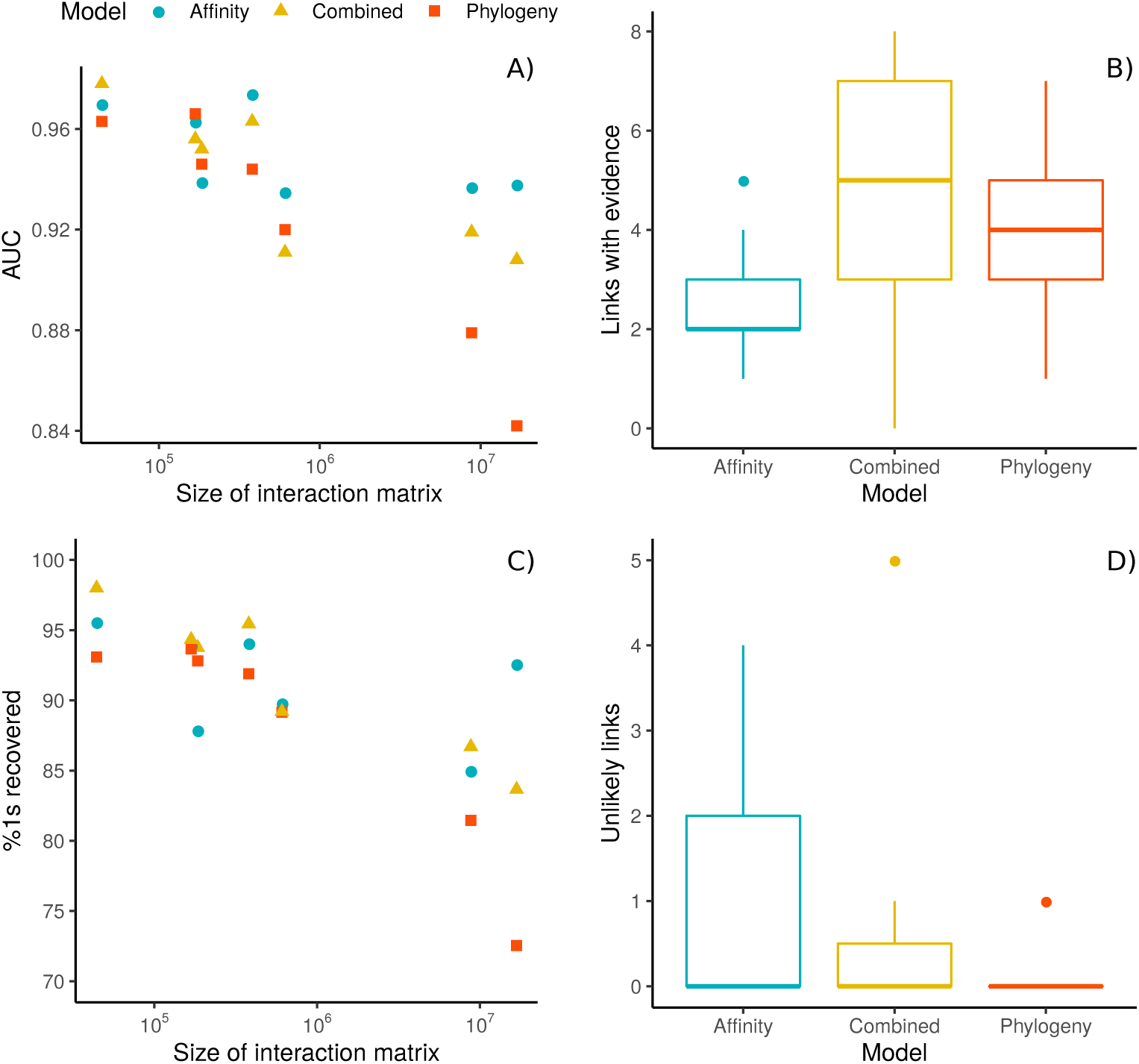
Model diagnostic plots, including internal predictive performance after 10-fold cross-validation (A,C) (see SM 1.2 for details), and the results of targeted literature searches for the top 10 undocumented links per model (B,D). Panel A shows area under the receiver operating characteristic curve (AUC) in relation to the size of interaction matrix (total number of potential host-parasite links). Panel C shows the percent documented interactions (1s) correctly recovered from the held-out portion in relation to the size of interaction matrix. Panels B and D show boxplots of the number of links with published evidence external to the original dataset (B), and the number of unlikely links based on ecological mismatch (D) out of the top ten most probable yet undocumented interactions for each of the three model types (affinity, combined, and phylogeny) run across the full dataset and each of the models subset by parasite type (Arthropods, Bacteria, Fungi, Helminths, Protozoa, Viruses).

As would be expected, the affinity only model tended to predict links between species with many previously documented associations, but this behavious was decreased in the combined model, and largely absent in the phylogeny only model (Fig. 3, SM 2.2). To further explore how each of the link prediction models propagated potential research biases, we compared the predicted probabilities per interaction to the degree product (calculated per host-parasite interaction by multiplying the observed host degree by the observed parasite degree). The degree product represents the basic expectation of a given interaction based on host and parasite affinities, and a measure of research effort per interaction. Comparing the observed degree products to the predicted interaction probabilities we can see that the predictions from the affinity only model are highly corellated with the observed host and parasite degrees (r=0.95; Fig. SM 3a), but again the correlation is reduced with the combined model (r=0.82; Fig. SM 3b), and further reduced with the phylogeny only model (r=0.49; Fig. SM 3c). This implies that predictions from the affinity model (and, to a lesser extent, the combined model) may be more likely to capture the signature of study effort when compared to the phylogeny model.

**Figure 3:**
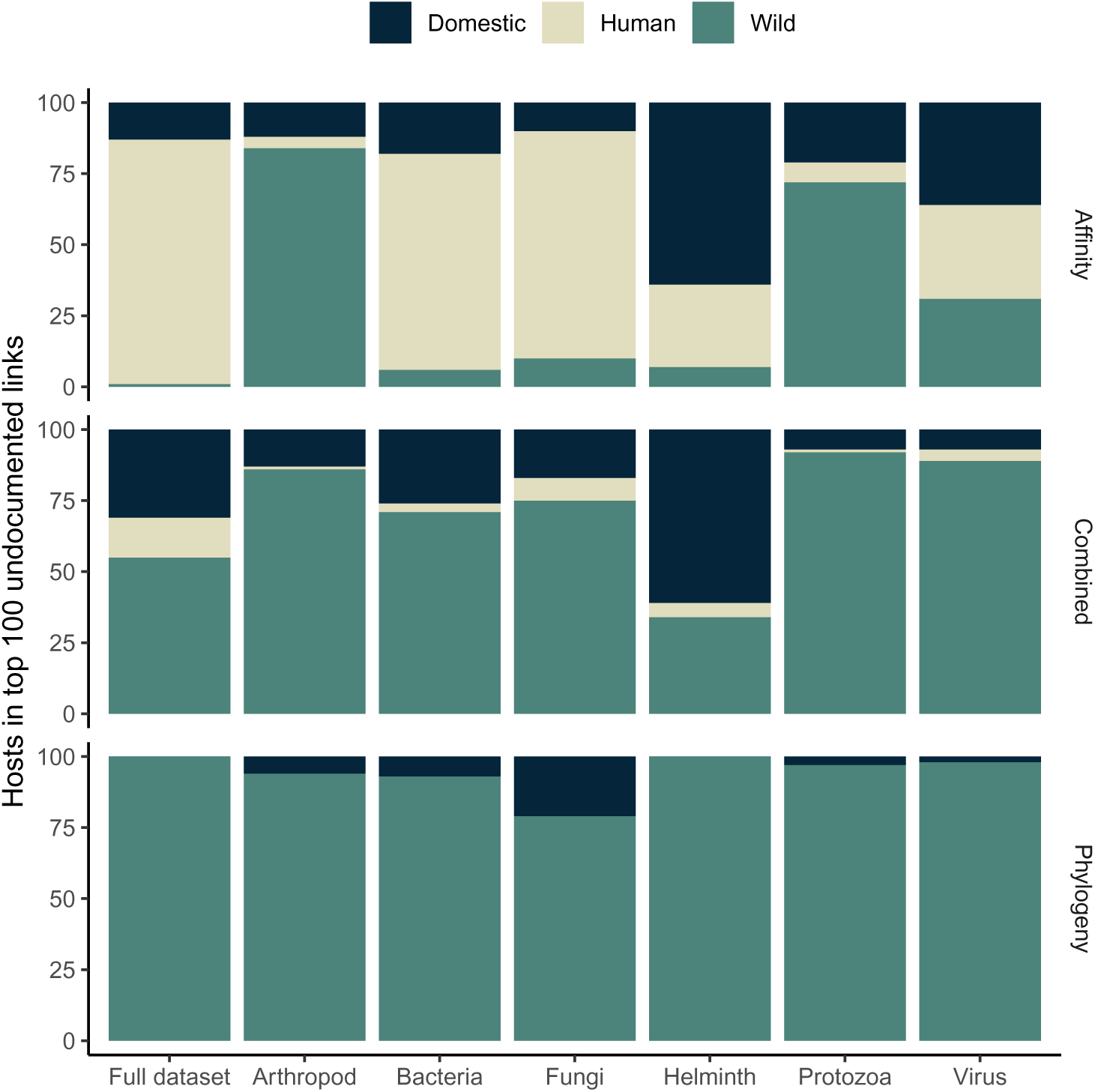
The frequencies of host types (human, domestic, and wildlife) included in the top 100 predicted links per model and dataset combination, which were not documented in the original database. Plots are grouped by data subset as columns (the full dataset, and subsets by parasite type), and model as rows (affinity, combined, and phylogeny).

Hotspot maps of undocumented host-parasite interactions (Fig. 4) illustrate large differences in the spatial distribution of missing links. Predictions from the affinity model (Fig. 4A) highlight Europe as a hotspot, while those from the phylogeny model reveal highest density of missing links predicted in tropical and central America, followed by tropical Africa and Asia (Fig. 4B). The hotspot map generated by the combined model was closer to that produced by the affinity only model, but with higher relative risk in sub-Saharan Africa, south America, and parts of southeast Asia (Fig. 4C).

**Figure 4:**
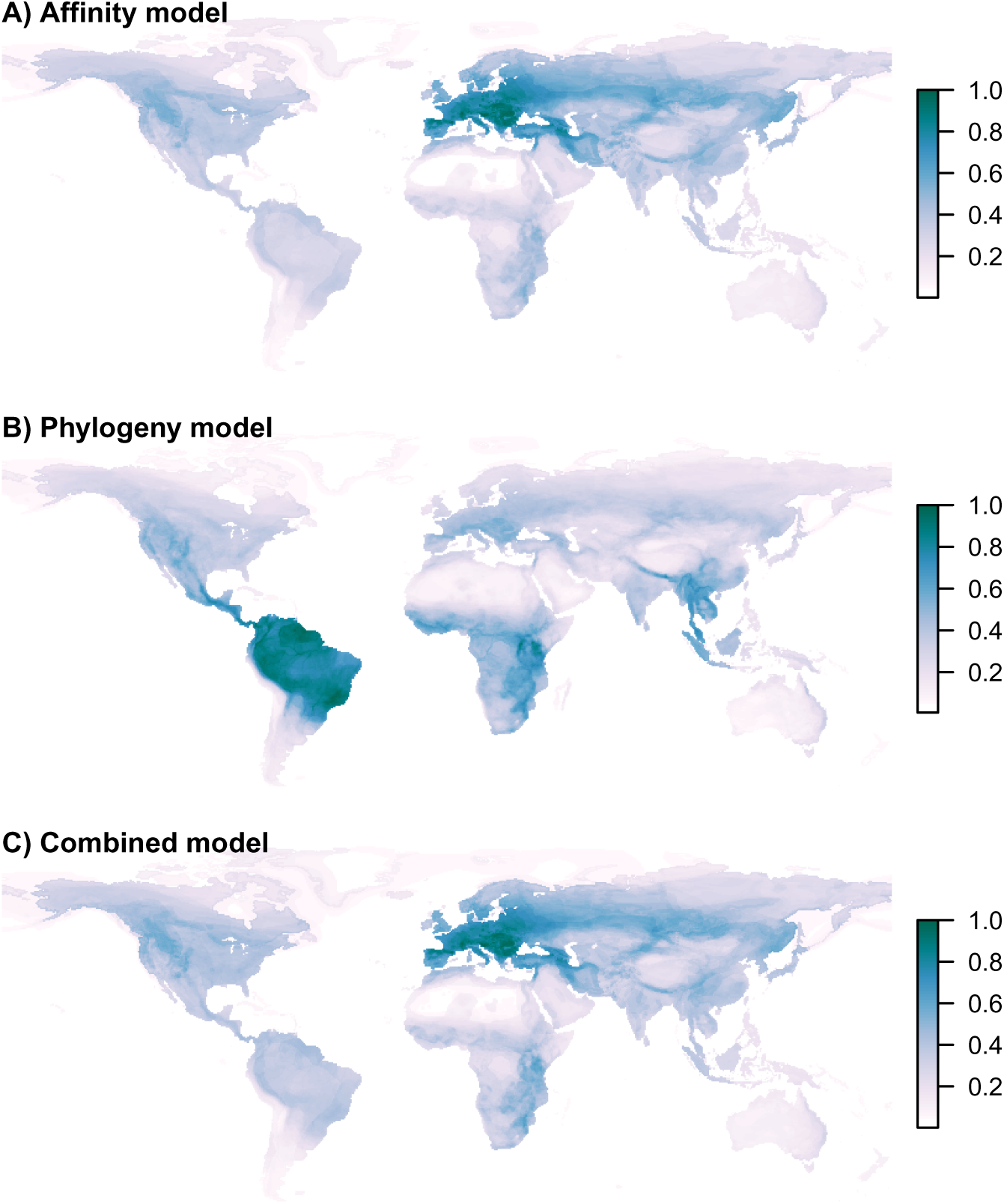
Global hotspot maps showing the relative density of predicted but undocumented host-parasite interactions. Maps were generated by summing the probabilities of undocumented host-parasite interactions per host species, summing across host species per cell (0.5 degree resolution) with range maps for terrestrial mammals from the IUCN, then standardizing to a scale of 0 - 1. A) represents the relative undocumented link density based on predictions from the affinity model, while B) represents the phylogeny model, and C) the combined model.

The top ranked links from the affinity and combined models were largely dominated by humans and domesticated animal hosts, while the phylogeny models more often predicted links among wildlife (Fig. 3) including endangered and relatively poorly studied species, some of which are critically endangered (IUCN, 2019). Parasites infecting large numbers and phylogenetic ranges of hosts were most often included in the top undocumented links in all models (ex. *Rabies lyssavirus*, *Sarcoptes scabiei*, *Toxoplasma gondii*, *Trypanosoma cruzi*). These parasites are commonly cited as capable of causing disease in a large number of (and sometimes all) mammals, though the majority of which have not been directly investigated. The combined model included a larger diversity of parasite species among the top predicted links (Table SM 2).

By conducting targeted literature searches of the top predicted missing links for each model by data subset, we found multiple links had published support, but were not included in the original source databases (See SM for lists of top links and detailed results of literature searches). The combined model identified a greater number of links with literature support but which were not in the original database (46/90) compared to the phylogeny (39/90) and affinity (29/90) models (Table SM 2).

## Discussion

Using a phylogeny-informed bipartite network model we were able to accurately predict missing links in a very large global mammal-parasite network. That we are able to make robust predictions even with extremely sparse input data, indicates that this modelling approach may be useful in other large, data-poor ecological networks. We compared the performance of our joint model with an affinity only (“rich-get-richer”) model, and a phylogeny only model. While predictive performance in cross-validation was regularly higher for the affinity model, our literature searches showed the phylogeny model to be less prone to predicting ecologically unlikely links, indicating that it may better capture the underlying ecological and evolutionary processes structuring host-parasite interactions. The layering of the affinity and phylogeny models allows the combined model to exploit the scale-free nature of many real-world networks while further correcting for ecologically unlikely links. The influence of this correction factor is best shown by contrasting the posterior interaction matrices (Fig. SM 2).

Predictions from the affinity model tended reflect existing degree distributions (SM 2.2), indicated by the dominance of humans and domesticated animals among the top predictions (Fig. 3, and highlighting Europe as a hotspot of undocumented links (Fig. 4), likely representing the volume of research on well-studied taxa in this region. We might expect that predicted links among well studied taxa would already be well-documented if common. However, affinity-based predictions may be critical for large-scale public health initiatives if they implicate widespread or abundant species as reservoirs of emerging infections. The affinity model is also more likely to identify parasite sharing among distantly related hosts, which has the potential to result in high mortality following host shifts (Farrell & Davies, 2019). While the affinitly model did predict links supported by our literature search, the top predictions included multiple links that are unlikely due to a mismatch in host ecology. For example, domestic cattle (*Bos taurus*) are predicted to be susceptible to infection by *Anisakis simplex*. *A. simplex* is a trophically transmitted nematode that uses aquatic mammals as final hosts, with marine invertebrates and fish as intermediate hosts (Buchmann & Mehrdana, 2016), implying that cattle may only be exposed to the parasite if fed a marine-based diet.

The phylogeny model uses only the evolutionary relationships among hosts to predict missing links, and was found to strongly mitigate the propagation of potential research biases compared to the affinity model (SM 2.2). Hotspots of predicted links from the phylogeny model show a spatial distribution in stark contrast to the affinity model, with highest density in tropical and central America, followed by tropical Africa and Asia (Fig. 4). It is unsurprising that these under-studied regions, with high host and parasite diversity, are identified as centres of undocumented host-parasite associations. The top undocumented links predicted by the phylogeny model also included fewer ecologically unlikely interactions, with only one unlikely interaction among those we investigated via literture review. *Echinococcus granulosus* is typically maintained by a domestic cycle of dogs eating raw livestock offal (Otero-Abad & Torgerson, 2013), and while wild canids such as *Lycalopex gymnocercus* are known hosts (Lucherini & Luengos Vidal, 2008), the phylogeny model predicts *Lycalopex vetulus* as a potential host. However, this interaction is unlikely as *L. vetulus* has a largely insectivorous diet (Dalponte, 2009), unlike the other members of its genus.

### Missing links

We identified a number of parasites which may impact the health of humans and domesticated animals. These include parasites currently considered a risk for zoonotic transmission such as *Alaria alata*, an intestinal parasite of wild canids – a concern as other *Alaria* species have been reported to cause fatal illness in humans (Murphy *et al*., 2012), and *Bovine viral diarrhea virus 1*, which is not currently considered to be a human pathogen, but is highly mutable, has the ability to replicate in human cell lines, and has been isolated from humans on rare occasions (Walz *et al*., 2010). However, there is a large amount of effort that goes into studying infectious diseases of humans and domestic species, and it is likely that most contemporary associations among humans and described parasites have been recorded, even if not included in the aggregated databases because they occur rarely or are difficult to detect. For example, we predicted that humans could be infected by *Bartonella grahamii*, and found that the first recorded case was in an immunocompromised patient in 2013 (Oksi *et al*., 2013). Similarly, humans are predicted to be susceptible to *Mycoplasma haemofelis*, which was again reported in someone who was immunocompromised (dos Santos *et al*., 2008), indicating that while these infections may pose little risk for a large portion of the human population, they are a serious concern for the health of immunocompromised individuals. These examples demonstrate our framework has the capacity to predict known human diseases, highlight parasites that are recognized zoonotic risks, and identify a number of parasites that are currently unrecognized as zoonotic risks.

Applying link prediction methods to wildlife global host-parasite networks can highlight both historical and contemporary disease threats to biodiversity, and identify parasites with the potential to drive endangered species towards extinction. For example, the phylogeny models identified links reported only in literature from over 30 years ago, such as *T. cruzi* in the critically endangered cotton-top tamarin (*Saguinus oedipus*) (Marinkelle, 1982) and the vulnerable black-crowned Central American squirrel monkey (*Saimiri oerstedii*) (Sousa, 1972). Our guided literature search also found evidence of severe infections in several endangered species such as rabies and sarcoptic mange (*Sarcoptes scabiei*) in Dhole (*Cuon alpinus*) (Durbin *et al*., 2005) and *Toxoplasma gondii* in critically endangered African wild dogs (*Lycaon pictus*) which caused a fatal infection in a pup (Van Heerden *et al*., 1995). Our model also predicts that rabies and sarcoptic mange are likely to infect the endangered Darwin’s fox (*Lycalopex fulvipes*). Disease spread via contact with domestic dogs (notably *Canine distemper virus*) is currently one of the main threats to this species (Silva-Rodŕıguez *et al*., 2016). Considering that both rabies and sarcoptic mange from domestic dogs are implicated in the declines of other wild canids, they may pose a serious risk for the conservation of Darwin’s foxes.

Because we did not include geographic constraints in our model, we may predict interactions among potentially compatible hosts and parasites that may be unrealized in nature due to lack of geographic overlap. These included *T. cruzi*, which is currently restricted to the Americas (Browne *et al*., 2017), infecting endangered African species such as black rhinoceros (*Diceros bicornis*), lowland gorilla (*Gorilla gorilla*), and chimpanzee (*Pan troglodytes*). Although contemporary natural infections of chimpanzees by this parasite are unlikely due to geography, we found a report of a fatal infection of a captive individual in Texas (Bommineni *et al*., 2009). In addition to this example, our models identified multiple infections documented only in captive animals. While we found no published cases of natural infections, these demonstrate that the model is able to identify biologically plausible infection risks that are relevant for captive populations, and may present future risks in the face of host or parasite translocation.

### Iterative link prediction and parasite surveillance

Link prediction in host-parasite networks is a critical step in an iterative process of prediction and verification, whereby likely links are identified, queried, and new links are added, allowing predictions to be updated. For example, we identified a number of interactions that were first documented in the literature only after the source databases were assembled (e.g. *Nematodirus spathiger* in *Gazella leptoceros* (Said *et al*., 2018), *Toxoplasma gondii* in *Papio anubis* (Kamau *et al*., 2016), *Trypanosoma cruzi* in horses (Bryan *et al*., 2016)), suggesting that prediction-guided literature searches offer a cost-effective solution to addressing knowledge gaps, maximizing the value of published literature, and identifying targets for future field-based sampling. We view our effort as working towards the larger goal of expanding and filling out the global mammal-parasite network, which also includes programs for pathogen discovery, and field and collections based parasite sampling.

We demonstrate that missing links in global databases of host-parasite interactions can be accurately identified using information on known associations and the evolutionary relationships among host species. Link prediction represents a cost-effective approach for augmenting global databases used in the study of disease ecology and evolution. As we move down the list of most probable links, we will uncover links with infectious organisms that are less well studied, but which may emerge as public or wildlife health burdens in the future. Through targeting research and surveillance efforts towards likely undocumented interactions, we can more efficiently gather baseline knowledge of the diversity of host-parasite interactions, ultimately supporting the development of fundamental theory in disease ecology and evolution (Stephens *et al*., 2016) and strengthening our understanding of disease spread and persistence in multi-host systems (Viana *et al*., 2014). In addition, prediction of host species that are susceptibile or exposed to known infectious diseases, we can guide the proactive surveillance of multi-host parasites underlying contemporary disease burdens, and those which may emerge in the future.

### Conclusion & Future Directions

In our study, all three models predicted links that were supported through targeted literature searches, though the host and parasite taxa differed. We therefore suggest that information on affinities and phylogeny are complementary, and while we place greater emphasis on the combined model here, choice of one model over another should depend on the goals of subsequent analyses, and whether it is important to minimize false positives or false negatives among predicted links. The phylogeny model leverages evolutionary relationships among taxa to generate predictions, however, the flexibility of the method allows for any information to be included, provided it can be represented as a distance matrix (Elmasri *et al*., 2020). Future models may be expanded to use trait or geographic distances among hosts to exclude ecological mismatches not currently captured by host phylogeny. Similarly, the model may be extended to incorporate weighted rather than binary associations, allowing for modelling links as a function of prevalence, intensity of infection, or to explicitly incorporate the amount of evidence supporting each link. In this way sampling intensity may be directly incorporated into link predictions, and help identify weakly supported interactions or sampling artefacts that may benefit from additional investigation.

We present our amalgamated interaction matrix and predictions of top missing links here as a resource for future work on the macroecology of multi host-multi parasite dynamics. However, important next step is to move beyond binary associations, and quantify the nature of the association between host and parasite (Becker *et al*., 2020). In this way we may be able to predict not only the presence or absence of a particular host-parasite interaction, but the epidemiological role each host plays in parasite transmission, the impact of infection on host fitness (Farrell & Davies, 2019), and better understand the ecologies of reservoir versus spillover hosts.

### Data & code

The amalgamated host-parasite matrix, results of all models, top predictions from each model, and R scripts to format predictions and reproduce the figures can be found at doi: 10.6084/m9.figshare.8969882. Code used to run the model is available at github.com/melmasri/HP-prediction.

## Acknowledgements

This work would not exist without the McGill Statistics-Biology Exchange (S-BEX) organized by Zofia Taranu, Amanda Winegardner, and Russel Steele. Additional thanks are due to Will Pearse, Ignacio Morales-Castilla, and Klaus Schliep for support and advice, and to Colin Carlson and Greg Albery for friendly review of the manuscript. MJF was supported by a Vanier NSERC CGS, the CIHR Systems Biology Training Program, the Quebec Centre for Biodiversity Science, and the McGill Biology Department. ME was supported by the McGill Department of Mathematics and Statistics, FRQNT, and an NSERC PDF.

## Author Contributions

MF and JD designed the study, ME, MF, JD, and DS designed the methods, MF compiled the data, MF and ME conducted the analyses, MF wrote the manuscript with input from JD. The authors declare no conflicts of interest.

## 1 Supplementary Methods

### 1.1 Database amalgamation

To amalgamate the source databases, host names were standardized to those in Wilson & Reeder (*1*) and used in the Fritz et al. (*2*) mammal phylogeny. Virus names were standardized to the 2015 version of the International Convention on Viral Taxonomy (*3*). For non-viral parasites there exists no single accepted authority for nomenclature or species concepts, however all parasite names were thoroughly checked for typographical and formatting errors. When potentially synonymous names were identified (e.g. *Cylicocyclus insigne* and *Cylicocyclus insignis*), online searches of primary literature and taxonomic databases including the Integrated Taxonomic Information System (www.itis.gov), Catalogue of Life (www.catalogueoflife.org), World Register of Marine Species (www.marinespecies.org), Encyclopedia of Life (www.eol.org), NCBI Taxonomy Database (www.ncbi.nlm.nih.gov/taxonomy), UniProt (www.uniprot.org), and the Global Names Resolver (resolver.globalnames.org) were conducted to resolve taxonomic conflicts. Synonymous names were corrected to the name with the majority of references, or to the preferred name in recently published literature or taxonomic revision when this information was available. Host associations for parasite names that were later split into multiple species were removed from the dataset (e.g. *Bovine papillomavirus*). All hosts and parasites reported below the species level were assigned to their respective species (e.g. *Alcelaphus buselaphus jacksoni* was truncated to *Alcelaphus buselaphus*), and any species reported only to genus level or higher (e.g. *Trichostrongylus sp.*) was removed. Our requirement for source databases to report Latin binomials facilitated taxonomy harmonization, but resulted in the exclusion of databases such as the GIDEON, which is a useful resource for modelling outbreaks of human infections based on disease common names, but does not provide unambiguous parasite scientific names (*4*). Of the 215 human diseases reported in Smith et al. (2014), we identified 197 that could be attributed to a predominant causal agent or group of organisms, all of which were already represented in our more comprehensive dataset. The remaining 18 human diseases represented those with no known causes (e.g. Brainerd diarrhea, Kawasaki disease), and those regularly caused by a diverse range of pathogens (e.g. viral conjuntivitis, invasive fungal infection, tropical phagedenic ulcer).”

### 1.2 Parameter estimation and predictive performance

For parameter estimation we split a dataset into ten folds, and used the MCMC algorithm described in Elmasri et al. (*5*) to estimate model parameters across each of the ten folds. For each model we determined the number of iterations required for parameter convergence by visual inspection of parameter traceplots, auto-correlation plots, and effective sample size (see Elmasri et al. (*5*) for detailed discussion of convergence diagnostics). For each fold we generated a posterior interaction matrix by averaging 1000 sample posterior matrices, where each sample matrix is constructed by drawing parameters at random from the last 10000 MCMC samples.

To assess predictive performance, we employed cross-validation across each of the ten folds used for parameter estimation. For each new fold, we set a fraction of the observed interactions (1s) to unknowns (0s), and attempted to predict them using a model fit to the remaining interactions. Here we only held out links for which there was a minimum of two observed interactions as the model would not be able to recover interactions for parasites that infect a single host species. Predictive performance was quantified using the area under the receiver operating characteristic (ROC) curve, which is a popular measure of potential predictive ability for binary outcomes. For each fold, an ROC curve is obtained by thresholding the predictive probabilities of the posterior interaction matrix, then calculating the true positive and false positive rates compared to the hold out set. In this way the posterior interaction matrix is converted to a set of binary interactions at the threshold value that maximizes the area under the ROC curve (AUC). To calculate ROC curves for each model-dataset combination, we took the average ROC values across the 10 folds. However, because our study is motivated by the belief that some 0s in our data are actually unobserved 1s, AUC may not be the best metric for comparing model performance as it increases when models correctly recover observed 0s. Therefore, in addition to AUC, we also assess predictive performance based on the percent of 1s accurately recovered, following a similar procedure. For prediction and guiding of the targeted literature searches, we generate a single posterior predictive interaction matrix for each model-dataset combination, by averaging these ten posterior interaction matrices.

## 2 Supplementary Results

**Figure SM 1:**
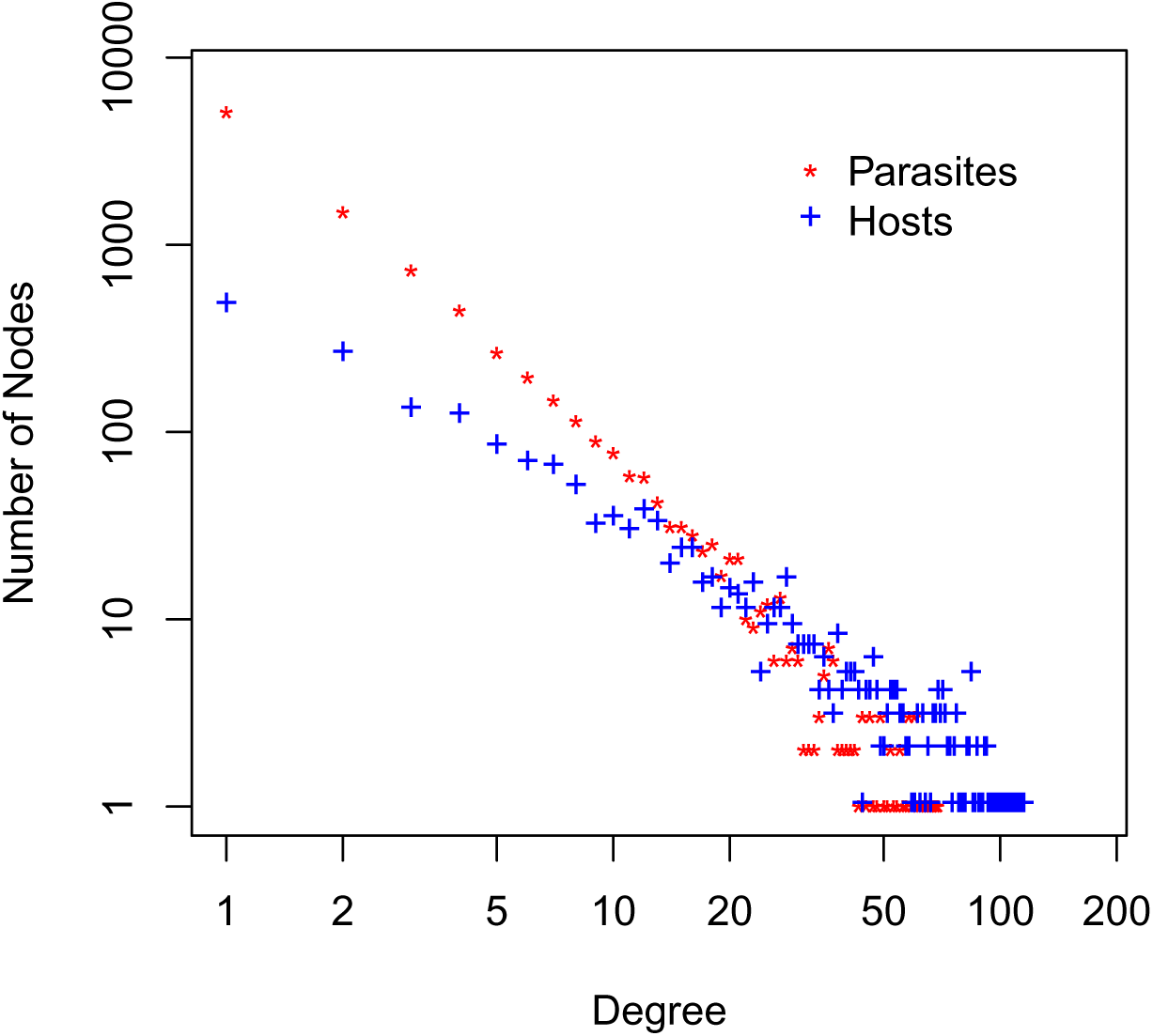
Degree distributions of the number of associations (degree) for hosts and parasites (nodes). The distributions are linear on the log scale, indicating a power-law in the number of observed associations per species across the network.

**Figure SM 2:**
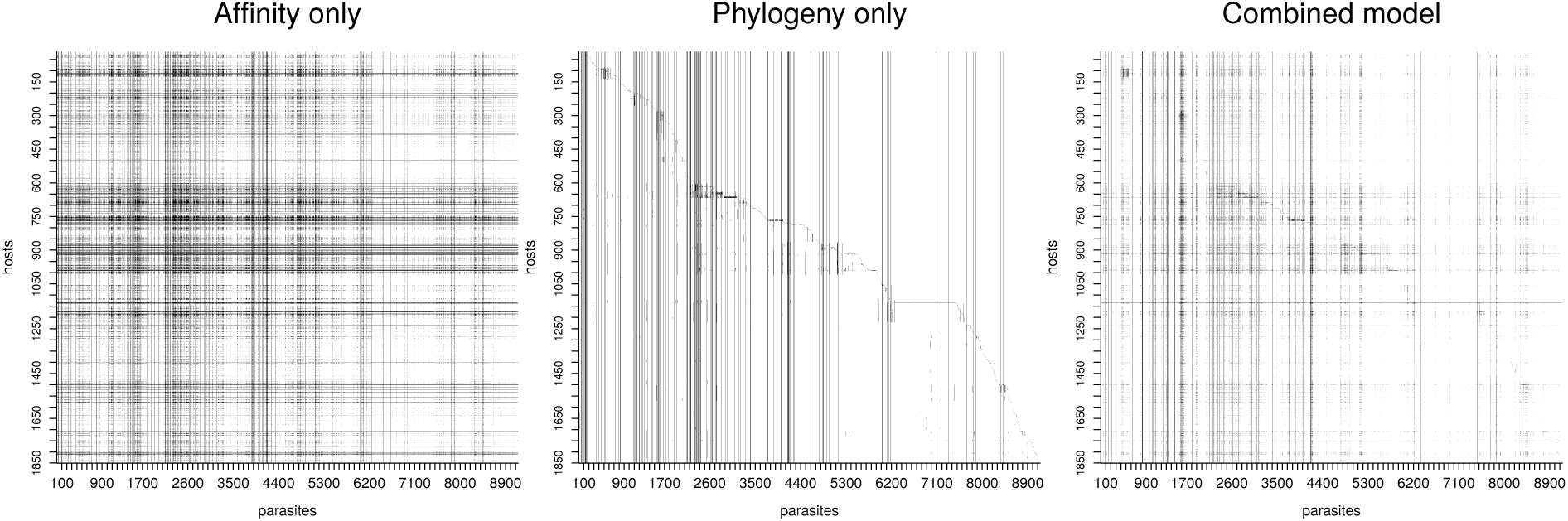
Posterior interaction matrices for the affinity, phylogeny, and combined models run on the full dataset. Axis labels represent indexes for hosts (rows) and parasites, with hosts ordered to match Figure 1.

### 2.1 Model diagnostics

All models showed high predictive accuracy in cross-fold validation: area under the receiver operating characteristic curve (AUC) values ranging from 0.842 - 0.978, where a maximum AUC of 1 signifies perfect predictive accuracy, and with between 72.54% and 98.00% of the held-out documented interactions successfully recovered (Fig. 2A, Table SM 1; see Fig. SM 2 for posterior interaction matrices for the full dataset). While remaining high overall, AUC tended to decrease with the total size of the interaction matrix, with the phylogeny only model demonstrating the lowest AUC when applied to the largest interaction matrices – the full matrix, and the helminth subset (Fig. 2A). The percent documented interactions (1s) correctly recovered from the held-out portion also decreased the total size of the interaction matrix (Fig. 2C), with the combined model outperforming the other models in all subsets except for the full dataset and the virus subset (Table SM 1).

This may reflect that successfully predicting held-out links becomes more difficult in more sparse matrices, which happens here as the size of the matrix increases. Further, for the phylogeny-only model, the larger drop in performance relative to other models may reflect variation in the optimal tree scaling parameter across different parasite subsets (see Figs. SM 9 & SM 9 for examples across parasite types). In the larger interaction matrices, additional parasite diversity is represented, implying that fitting a single tree transformation may result in sub-optimal prediction as the relative importance of shallow versus deep phylogenetic distances is likely to vary across parasite subgroups within these matrices. Future research may benefit from being able to fit multiple different scaling parameters per parasite group.

For each model, we conducted literature searches for the top ten most likely links without documentation in the current database. This resulted in 177 unique host-parasite links after removing duplicate predictions across models and subsets (see Materials & Methods). Of the undocumented links for which literature searches were conducted, we identified 72 links with evidence of infection (direct observation, genetic sequencing, or positive serology), and an additional 14 links with some evidence, but for which additional confirmatory data are required (e.g. antibodies but no confirmed cases for human infections, known cross-reactivity of the serological test used, an unconfirmed visual diagnosis, or the identification of a genetically similar but previously unknown parasite). Of the remaining links for which we could not find conclusive evidence, we highlight 39 that should be targeted for surveillance. These include links where there is known geographic overlap in the ranges of the host and parasite and host ecologies likely facilitate exposure. We also identify a number of links that are highly likely in the model, but are unlikely due to the mode of disease transmission, non-overlapping host and parasite geographies, or potential competitive interactions with closely related parasites. Overall the full and phylogeny only models tended to identify a greater number of links with published evidence, and fewer ecologically unlikely links (Fig. 2D).

**Table SM 1:**
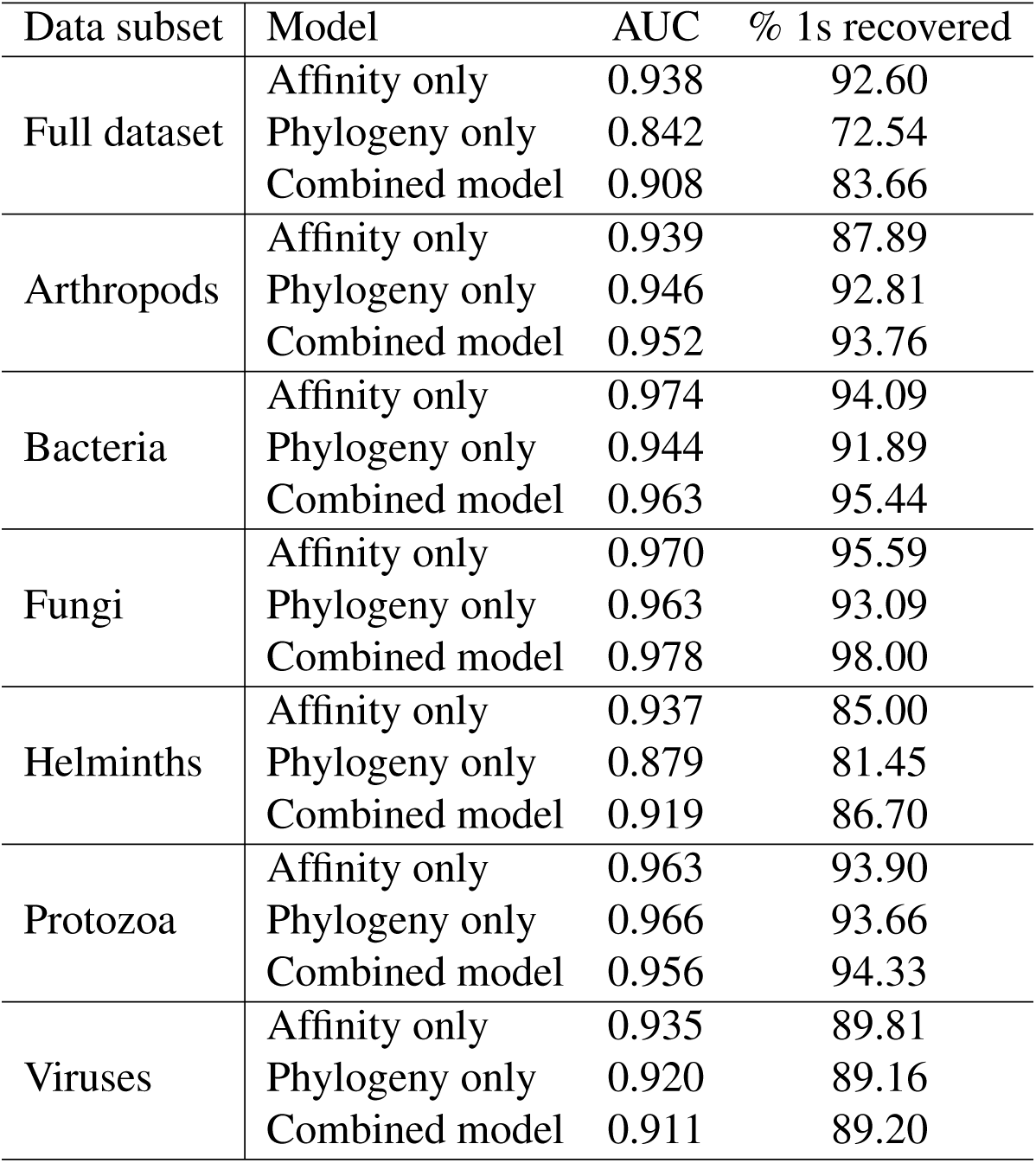
Average model performances diagnostics after 10-fold cross validation: area under the receiver operating characteristic curve (AUC), and percent documented interactions (1s) correctly recovered from the held-out portion.

### 2.2 Investigation of bias propagation

**Figure SM 3:**
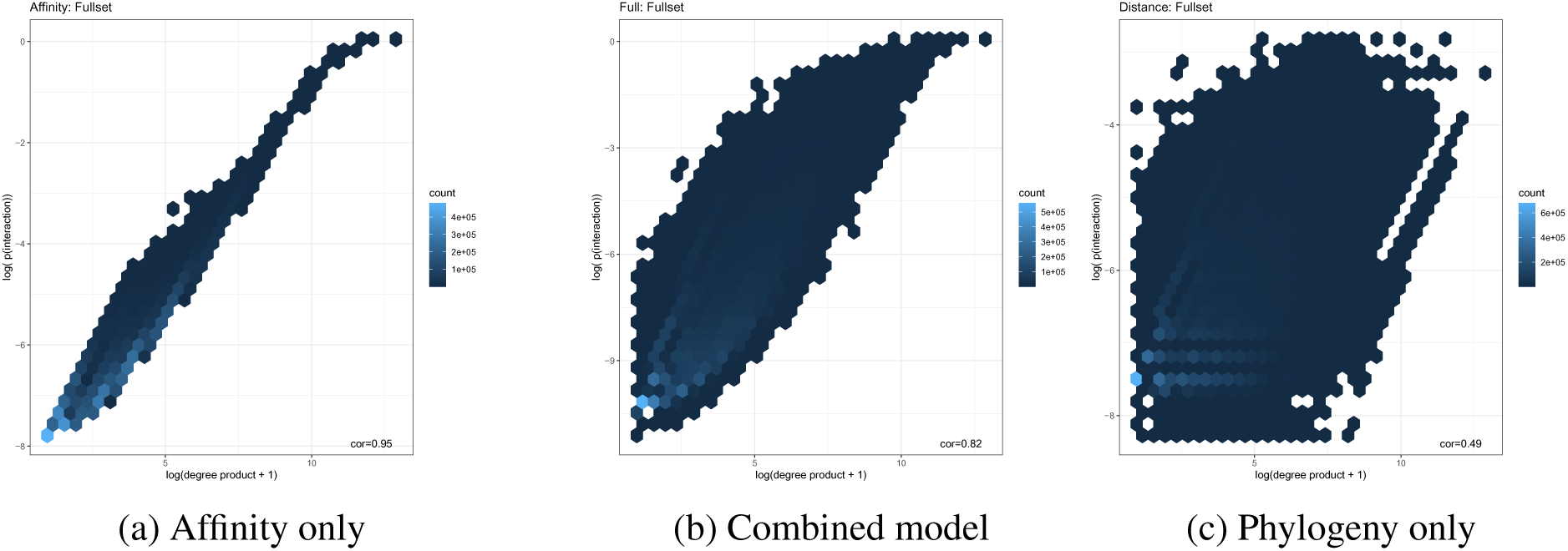
Hexbin plots of the relationship between the degree product per interaction (calculated as host degree multiplied by parasite degree) on the x axis (log+1 transformed), and the predicted probabiltiy based on a given model on the y axis (log transformed). Plots A-C represent the affinity, combined, and phylogeny models run on the full dataset. Single host parasites are removed prior to plotting.

**Figure SM 4:**
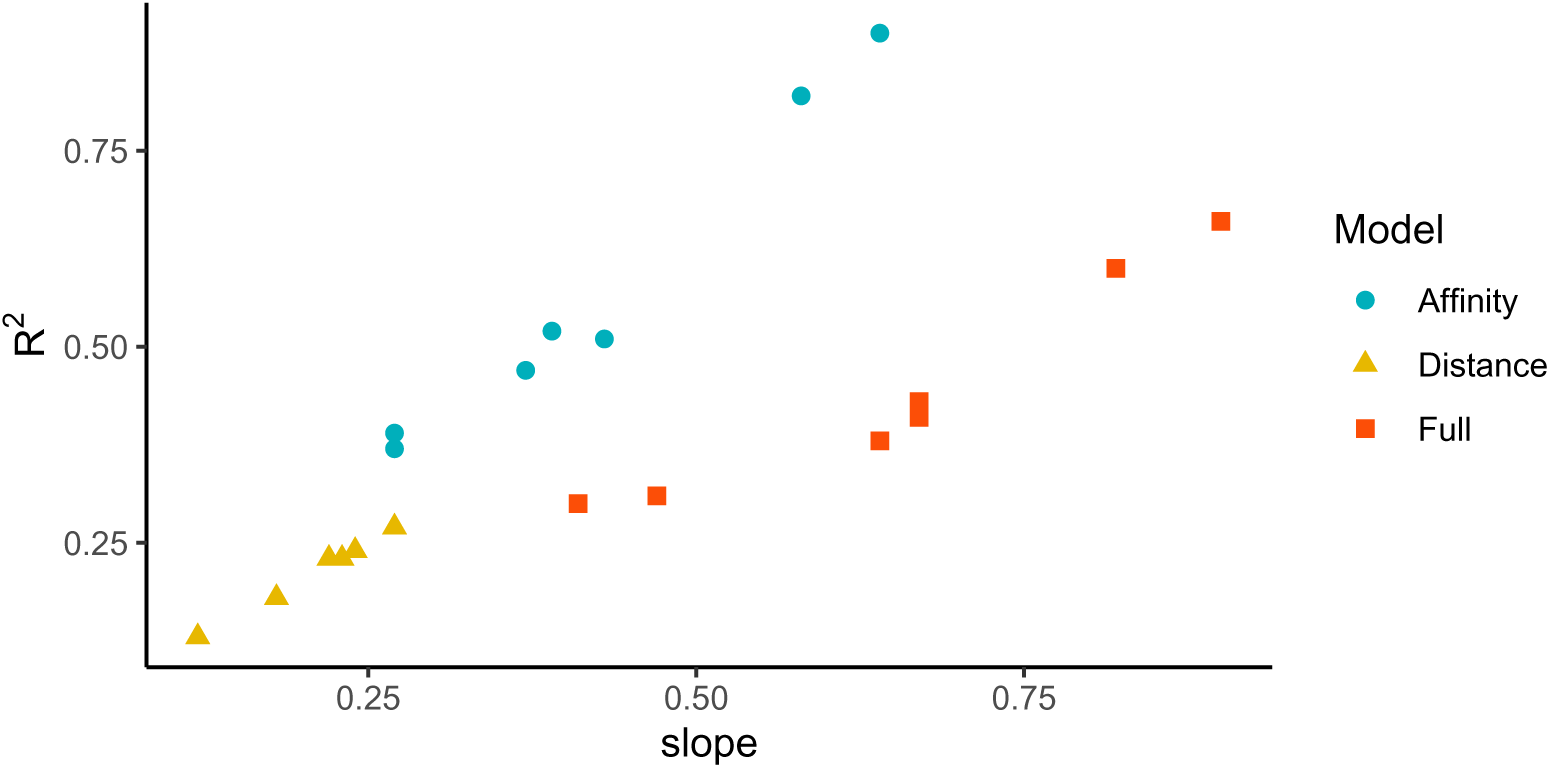
For each model-dataset combination, a linear regression relating degree product to predicted probabilty of each interaction was fit. The figure indicates the estimated slope and explained variance per linear regression.

**Figure SM 5:**
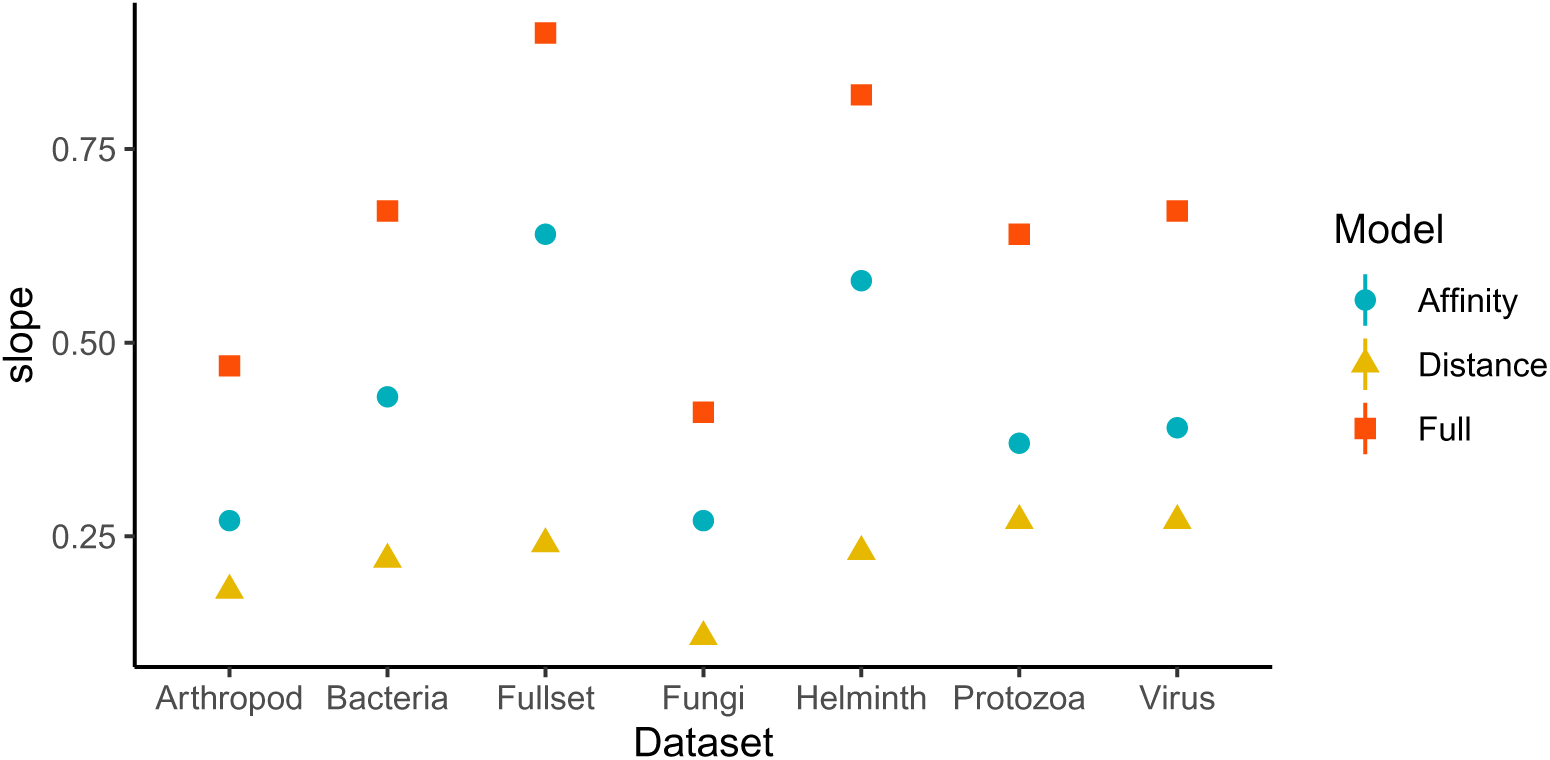
For each model-dataset combination, a linear regression relating degree product to predicted probabilty of each interaction was fit. The figure indicates the estimated slope between degree product and predicted probability. Note that 95% confidence intervals are plotted but too small to be seen.

**Figure SM 6:**
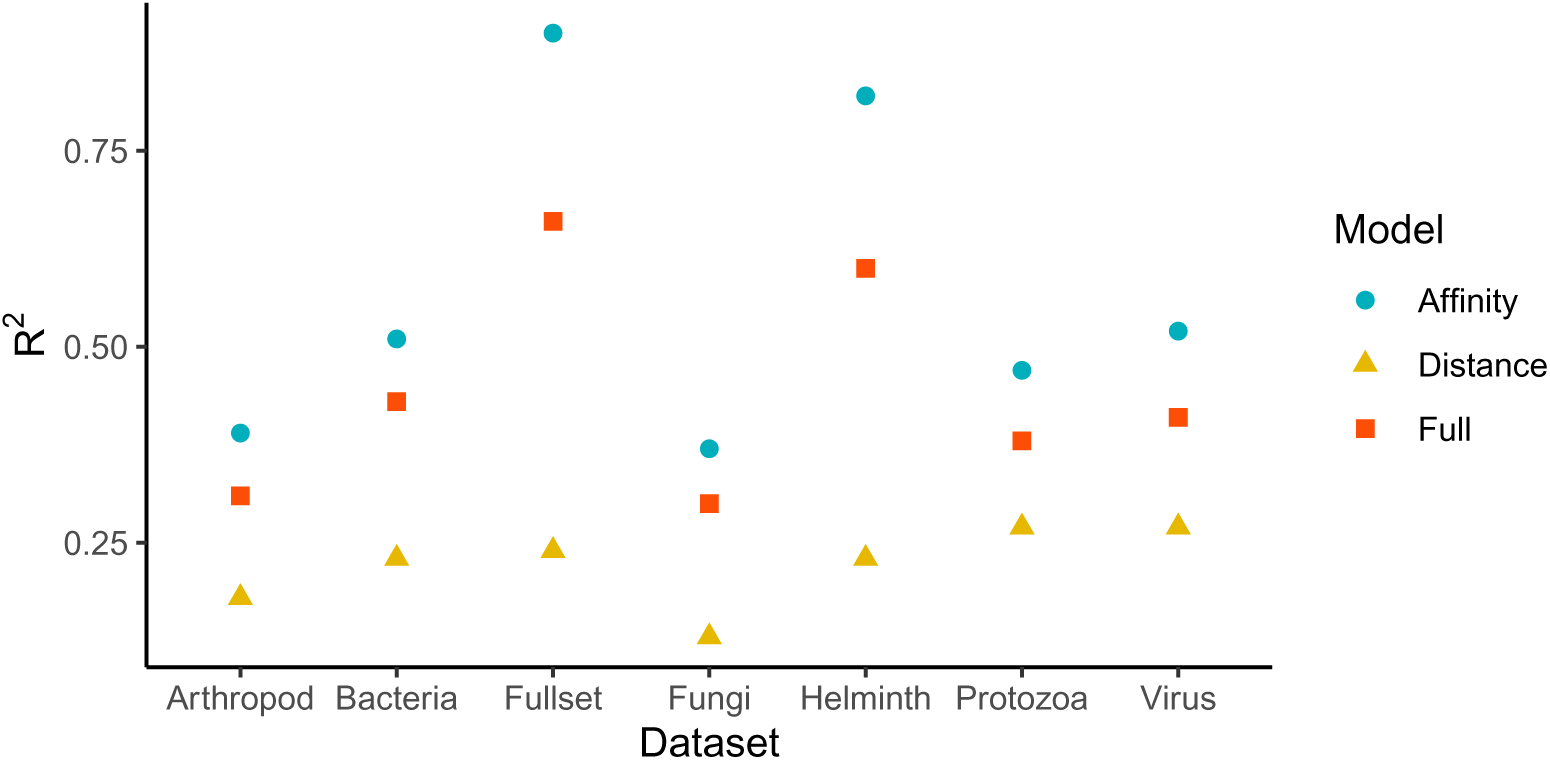
For each model-dataset combination, a linear regression relating degree product to predicted probabilty of each interaction was fit. The figure indicates the variance explained by degree product.

**Figure SM 7:**
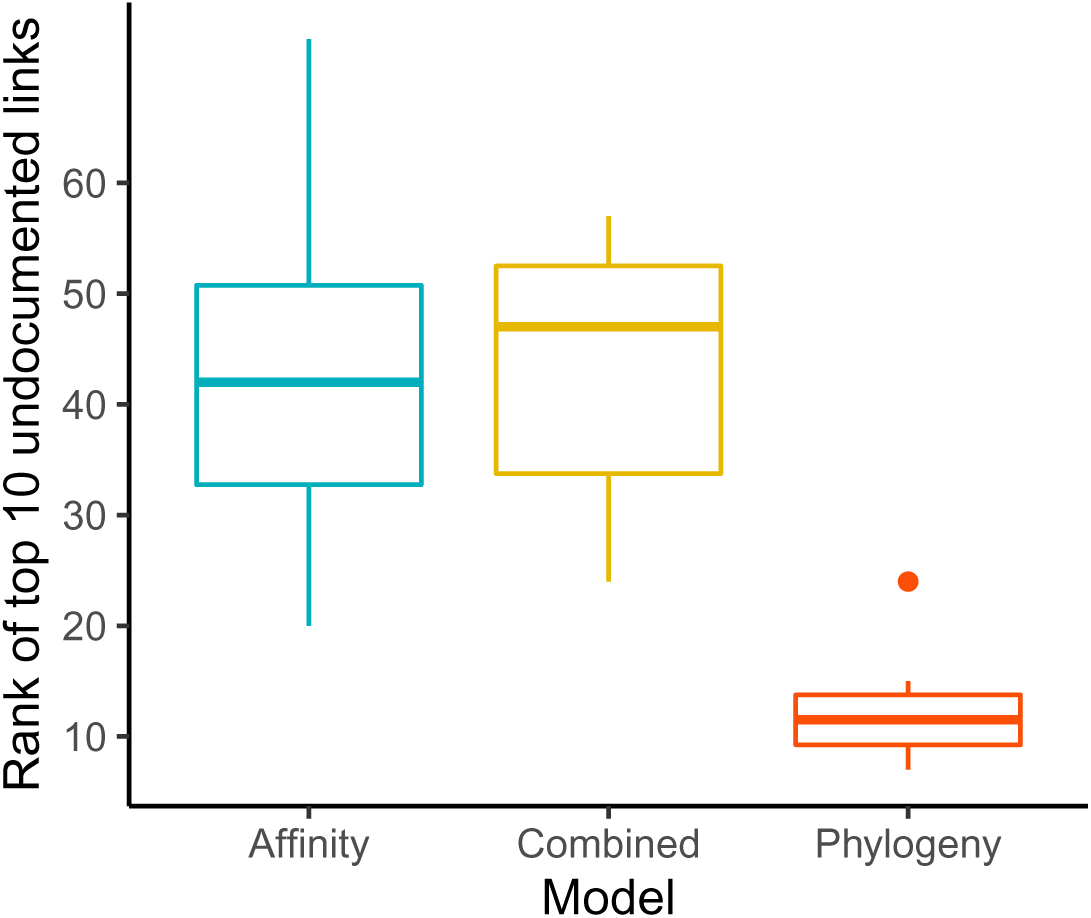
Boxplots of the ranks of the top 10 predicted links with no documentation in the original dataset, for each of the three model variants applied to the full dataset.

**Figure SM 8:**
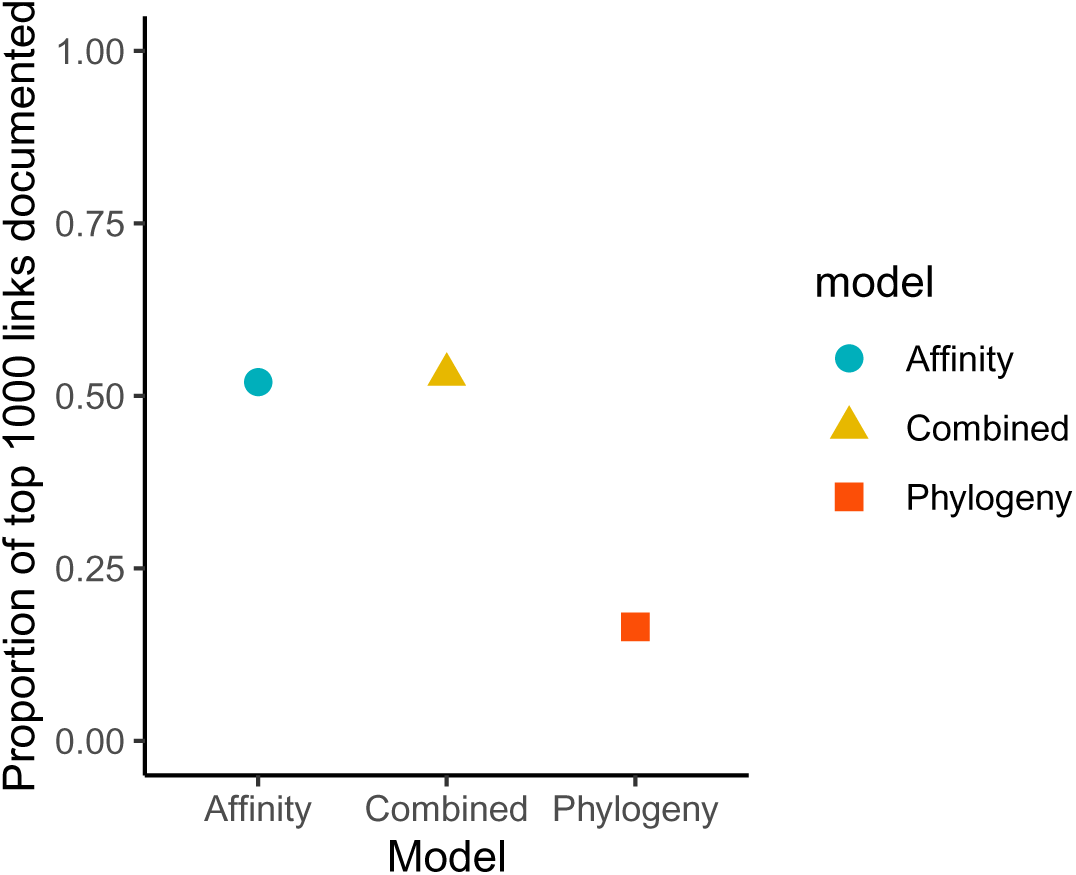
The proportion of the top 1000 predicted links which had already been documented in the original dataset. Results are shown for each of the model variants applied to the full dataset.

### 2.3 Phylogeny scaling

To account for uncertainty in the phylogenetic distances among hosts and improve prediction, the model estimates a tree scaling parameter (*η*) based on an early-burst model of evolution (*6*). Across models, *η* was estimated to be positive, corresponding to a model of accelerating evolution, and suggesting less phylogenetic conservatism in host link associations among closely related taxa than predicted under a pure-Brownian motion model (*6*). Not surprisingly, *η* varied when the data was subset by parasite type (Figs. SM 9 & SM 10). Interestingly, arthropods and fungi were estimated to have the smallest *η* parameters (both *∼* 8.15), perhaps reflecting the tendency for fungi to include opportunistic pathogens such as *Pneumocystis carinii* and *Chrysosporium parvum*. In contrast, larger *η* was estimated for helminths and viruses (10.28 and 9.54 respectively), consistent with the observation of Park et al. (*7*) that mean phylogenetic specificity is similar in these two groups, though viruses are more variable and contain more extreme specialist and generalist parasites. Overall the full dataset was estimated with an *η* parameter most similar to the helminth subset (10.76), reflecting the representation of helminths in the full dataset (roughly 64% of the observed host-parasite interactions). The discrepancy in phylogeny scaling across the Combined model and the subsets by parasite type likely contributes to explaining why the performance of the phylogeny only model is lower in the full dataset. Future extensions may benefit from developing a more flexible model to allow for an interaction between phylogenetic scaling and parasite taxonomy, rather than using a single scaling parameter.

**Figure SM 9:**
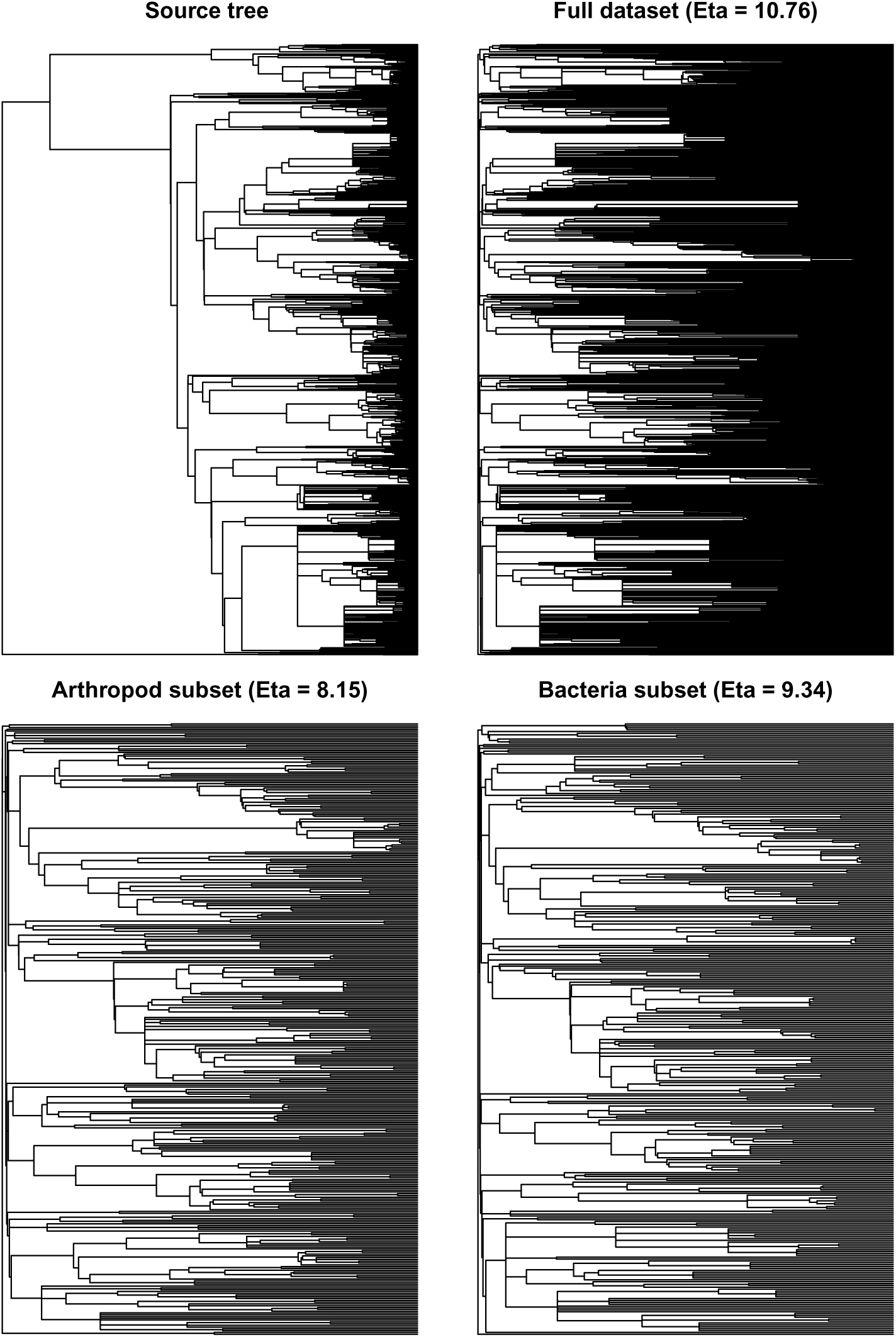
The source phylogeny pruned to include only hosts in the amalgamated dataset, and the phylogenies scaled by mean estimated Eta for the phylogeny only models applied to the full dataset, and arthropod and bacteria subsets.

**Figure SM 10:**
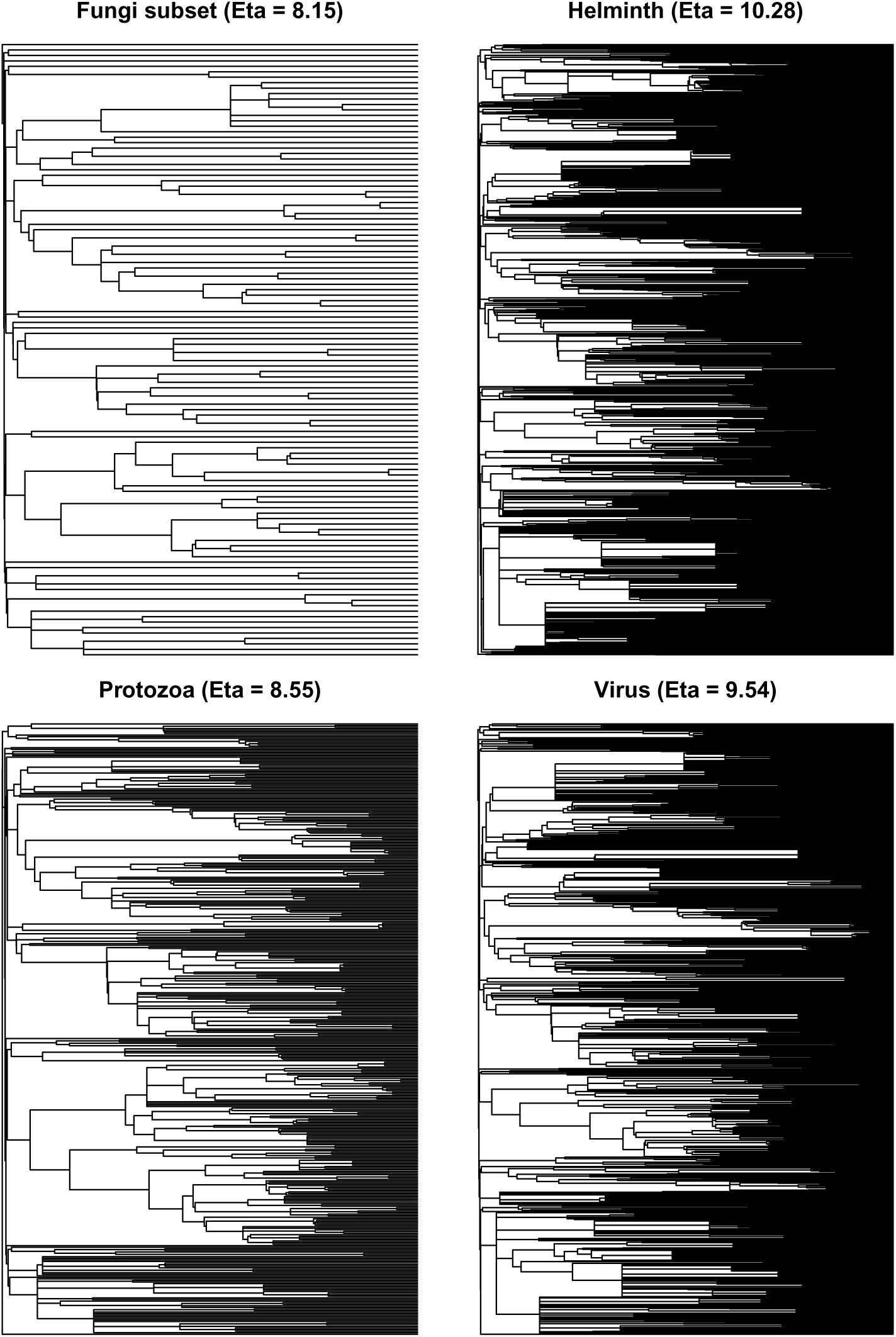
Phylogenies scaled by mean estimated Eta for the phylogeny only models applied to the fungi, helminth, protozoa, and virus subsets.

#### Top predicted links and literature search results

**Table SM 2:**
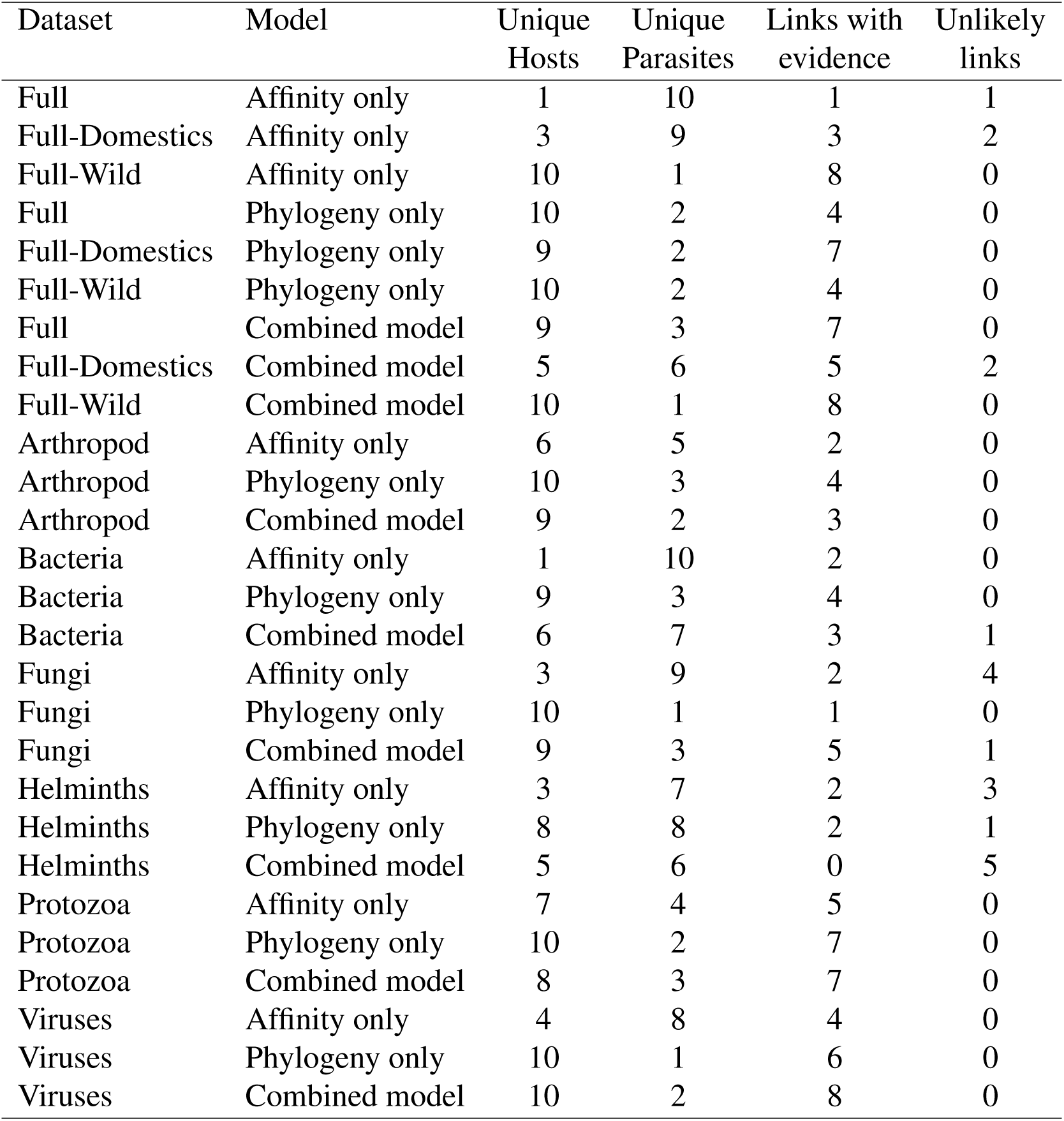
Numbers of unique hosts, unique parasites, links for which evidence was identified in the literature, and unlikely links based ecological mistmatch for each of the top 10 predicted links per model.

### 2.4 Affinity only – Full Dataset

**Table SM 3:** Top 10 undocumented links with highest probablity of interaction: Affinity only model - full data set.

| Host | Parasite | Status |
| --- | --- | --- |
| Homo sapiens | Taenia mustelae |  |
| Homo sapiens | Bluetongue virus |  |
| Homo sapiens | Bovine viral diarrhea virus 1 |  |
| Homo sapiens | Neospora caninum |  |
| Homo sapiens | Mastophorus muris |  |
| Homo sapiens | Plagiorchis vespertilionis | Confirmed |
| Homo sapiens | Carnivore protoparvovirus 1 |  |
| Homo sapiens | Alaria alata |  |
| Homo sapiens | Taenia pisiformis |  |
| Homo sapiens | Physocephalus sexalatus | Unlikely |

- A recent molecular phylogeny of the *Taenia* genus supported the creation of a new genus and renaming of *Taenia mustelae* to *Versteria mustelae* (*8*). Since then there has been a report of fatal infection of a previously unknown *Versteria* species in a captive orangutan *Pongo pygmaeus* cloesly related to species found in wild mustelids, suggesting the need for increased vigilance of *Versteria* infections in humans (*9*).
- While bluetongue virus is known to infect a wide range of ruminants, it is not currently considered to infect humans (*10*).
- Bovine viral diarrhea viruses are not considered to be human pathogens, but there is some concern about zoonotic potential as they are highly mutable, have the ability to replicate in human cell lines, and have been isolated from humans on rare occasions (*11*).
- Although antibodies to *Neospora caninum* have been reported in humans, the parasite has not been identified in human tissues and the zoonotic potential is not known (*12*).
- *Mastophorus muris* is a rodent-specific nematode that requires arthropods as intermediate hosts and while this makes it unlikely to infect humans, it was recently documented in an urban population of rats in the UK (*13*), indicating the potential for human exposure.
- While natural infections of *Plagiorchis* species in humans are rare, the first case of human infection by the bat parasite *Plagiorchis vespertilionis* was reported in 2007 (*14*). The source of infection is uncertain, it has been suggested that freshwater fish and snails may be undocumented intermediate hosts and infection was due to ingestion of raw freshwater fish.
- Carnivore protoparvovirus can infect a number of hosts in the order *Carnivora* (*15*), though there seems to be no evidence of human infection.
- *Alaria alata*, an intestinal parasite of wild canids, has not been identified in humans, but is considered a potential zoonotic risk as other *Alaria* species have been reported to cause fatal illness in humans (*16*).
- Due to the characteristics of the biological cycle of *Tenia pisiformis* and the observation that it is innocuous in humans, this parasite has been used as a model for the study of other important zoonosis relevant to human health including *T. solium* (*17*).
- The definitive hosts of *Physocephalus sexalatus* are commonly wild and domestic pigs, but it is sometimes found in other mammals and some reptiles (*18*). However the parasite uses beetles as an intermediate host, which makes human infection unlikely.

### 2.5 Affinity only – Full Dataset – Domestic Hosts

**Table SM 4:**
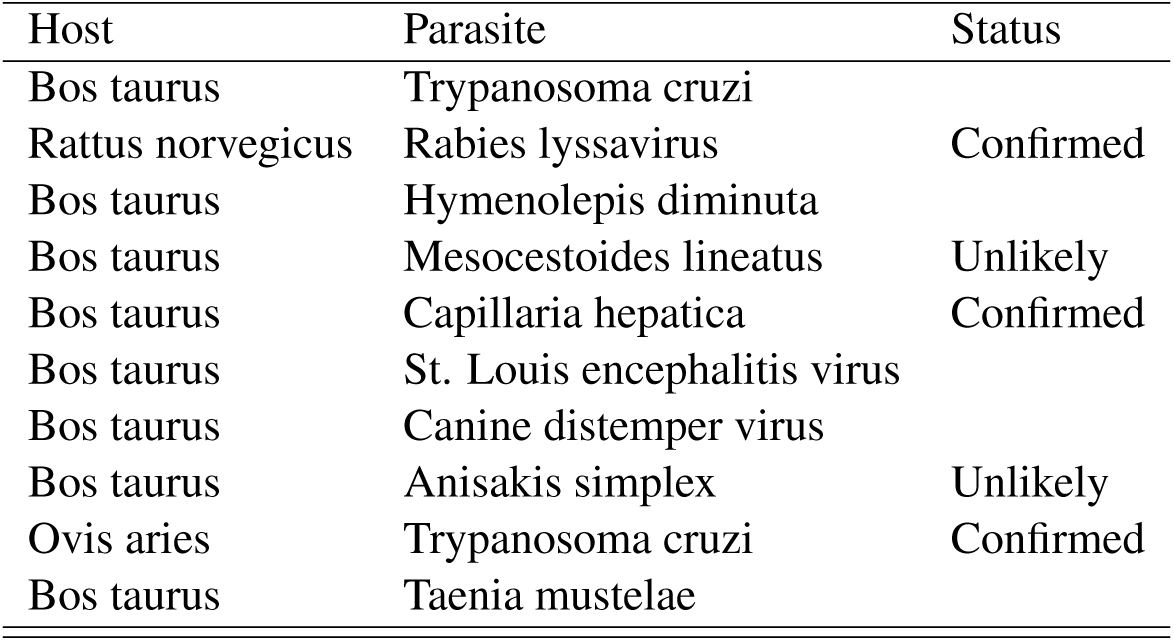
Top 10 undocumented links with highest probablity of interaction for domesticated species: Affinity only model - full data set.

- Currently the role of cattle in the epidemiology of Chagas disease (caused by *Trypanosoma cruzi*) is unknown, though the majority of cattle in Latin America may be exposed (280 million heads; 1/4 of the world population) (*19*). (*20*) report that 177 species have been documented as susceptible to infection by *T. cruzi*, with domestic hosts in some cases being responsible for the maintenance of local parasite populations over long periods of time. While cattle have tested positive in serological studies, cows and other domestic species are also infected by *Trypanosoma* and *Phytomonas* species which can cause cross-reactions in diagnostic tests (*21*).
- Rabies has an extremely large host range and surprisingly rats are rarely reported as suffering from rabies, though there have been a few reported cases rabid *Rattus norvegicus* in the United States (*22*).
- The natural hosts of *Hymenolepis diminuta* are rats and cattle have not been found to be susceptible to infection, however *H. diminuta* eggs have been found in the feces of dairy cattle, likely the result of ingesting forage contaminated with rodent feces (*23*).
- *Mesocestoides lineatus* has a three-stage lifecycle with two intermediate hosts and a large range of carnivorous mammals as definitive hosts (*24*). Human infections of the tapeworm *Mesocestoides lineatus* are rare but can occur through the consumption of chickens, snails, snakes, or frogs (*25*) and therefore it is unlikely that cows will ingest the intermediate life stages of this parasite.
- *Capillaria hepatica* (syn. *Calodium hepaticum*), is a globally distributed zoonotic parasite which uses rodents as main hosts, but is known to cause infection in over 180 mammalian species, including cattle (*26*).
- Birds are the primary vertebrate hosts for St. Louis encephalitis virus, though amplification by certain mammals has been suggested (*27*). There is some serological evidence of infection in domestic mammals, including cattle (*28*). The common vector *Culex nigripalpus* feeds primarily on birds, but shows a seasonal shift from avian hosts in the spring to mammalian hosts in the summer, indicating it may be able to act as a bridging vector among different host species (*27*).
- Canine distemper virus infects a wide range of hosts within the order Carnivora, but has also been found to cause fatal infection in some non-human primates and peccaries (*29*).
- *Anisakis simplex* uses cetaceans as final hosts, with marine invertebrates and fish as intermediate hosts (*30*). Whales are infected through ingestion, indicating that while cattle may be susceptible, though they would need sufficient exposure to marine based feed.
- *Ovis aries* is reported to be infected by *Trypanosoma cruzi* (*20*).
- We found no evidence of *Bos taurus* infection by *Taenia mustelae*.

### 2.6 Affinity only – Full Dataset – Wild Hosts

**Table SM 5:** Top 10 undocumented links with highest probablity of interaction for wild host species: Affinity only model - full data set.

| Host | Parasite |  |
| --- | --- | --- |
| *Cervus elaphus* | Rabies lyssavirus | Confirmed |
| *Ondatra zibethicus* | Rabies lyssavirus | Confirmed |
| *Capreolus capreolus* | Rabies lyssavirus | Confirmed |
| *Sorex araneus* | Rabies lyssavirus |  |
| *Apodemus sylvaticus* | Rabies lyssavirus | Confirmed |
| *Odocoileus virginianus* | Rabies lyssavirus | Confirmed |
| *Syncerus caffer* | Rabies lyssavirus |  |
| *Dama dama* | Rabies lyssavirus | Confirmed |
| *Pan troglodytes* | Rabies lyssavirus | Confirmed |
| *Microtus arvalis* | Rabies lyssavirus | Confirmed |

- For the affinity only model run on the full dataset, all of the top ten interactions involve rabies, which is unsurprising as rabies is commonly said to be capable of infecting all mammal species (*31*) and is the parasite with the largest number of documented hosts in the database (176). Of these ten most likely hosts, we found evidence of rabies infection in eight (*Cervus elaphus* and *Odocoileus virginianus* (*32*), *Ondatra zibethicus* (*33*), *Capreolus capreolus* (*34*), *Pan troglodytes* (*35*), *Apodemus sylvaticus* (*36*), *Dama dama* (*37*), and successful experimental infection of *Microtus arvalis* (*38*)).

### 2.7 Phylogeny only – Full Dataset

**Table SM 6:**
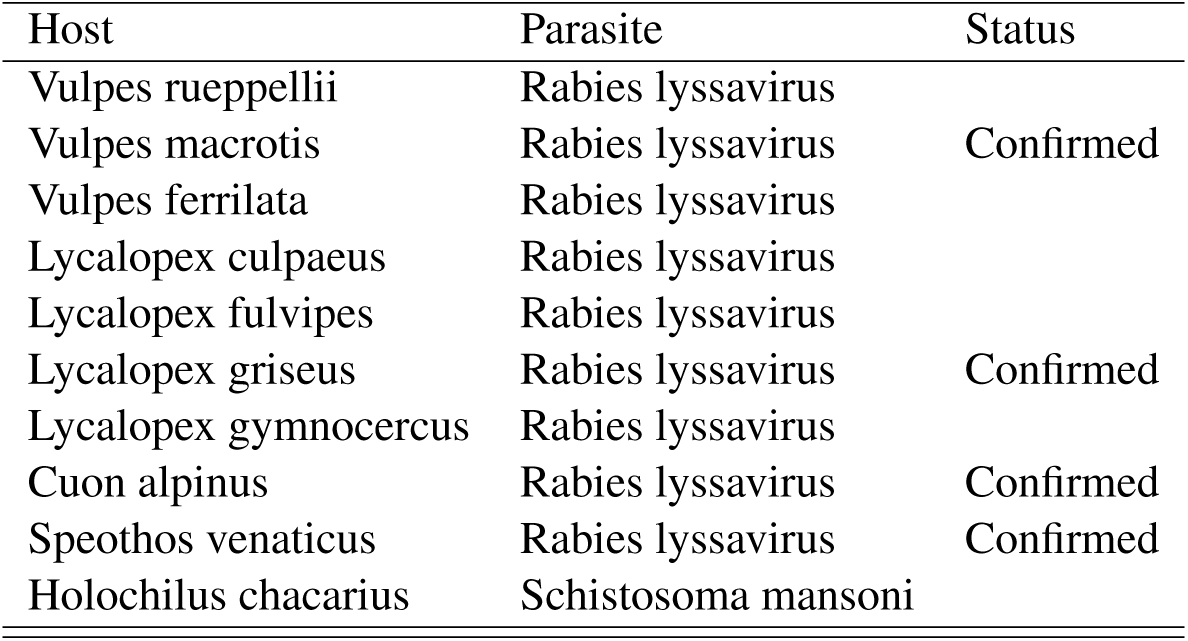
Top 10 undocumented links with highest probablity of interaction (Phylogeny only model - full data set)

- We did not find any record of rabies infecting *Vulpes rueppellii*, but this should be investigated as this disease is known to cause severe declines in wild canids (*39*).
- Rabies has been documented to infect the endangered San Joaquin kit fox (*Vulpes macrotis mutica*) and is suggested to have caused a catastrophic decline of the species in the 1990s (*40*).
- Domestic dogs are considered a major predation threat to the Tibetan fox (*Vulpes ferrilata*) (*41*), and rabies is confirmed to circulate in wild and domestic animals in Tibet (*42*).
- Domestic dogs alter the ecology of Andean foxes (*Lycalopex culpaeus*), have been observed hunting them, and are a potential source of infection (*43*).
- Diseases from domestic dogs (largely canine distemper) is considered a major threat to the endangered *Lycalopex fulvipes* (*44*), indicating that rabies may also pose a risk.
- There is one report of serological evidence of rabies in *Lycalopex griseus* in Chile in 1989 (*45*) (reported as *Pseudalopex griseus*, a formerly accepted name (*1*)).
- We did not find evidence of rabies infection in *Lycalopex gymnocercus*, however its distribution overlaps with species known to be important in the transmission of rabies in Brazil (*46*).
- The endangered Dhole (*Cuon alpinus*) is known to suffer from rabies and was a source of fatal human infections during an outbreak in the 1940s (*47*).
- Rabies has been reported as potentially infecting *Speothos venaticus* (*48*) and there is a report of an individual with positive serology (*49*).
- We could not find evidence of *Schistosoma mansoni* infection in *Holochilus chacarius*. Congener *Holochilus braziliensis* was experimentally shown to be a viable host, although infection resulted in host death (*50*).

### 2.8 Phylogeny only – Full Dataset – Domestic Hosts

**Table SM 7:** Top 10 undocumented links with highest probablity of interaction for domesticated species: Phylogeny only model - full data set.

| Host | Parasite | Status |
| --- | --- | --- |
| Bison bison | Rabies lyssavirus | Confirmed |
| Bos grunniens | Rabies lyssavirus | Confirmed |
| Bos frontalis | Rabies lyssavirus |  |
| Bos javanicus | Rabies lyssavirus |  |
| Vicugna vicugna | Rabies lyssavirus |  |
| Rattus rattus | Rabies lyssavirus | Confirmed |
| Rattus norvegicus | Rabies lyssavirus | Confirmed |
| Cavia porcellus | Rabies lyssavirus | Confirmed |
| Oryctolagus cuniculus | Rabies lyssavirus | Confirmed |
| Bos grunniens | Toxoplasma gondii | Confirmed |

- Rabies in *Bison bison* is considered rare, but there are multiple cases reported (*51*).
- Rabies has been reported to infect yak (*Bos grunniens*) in Nepal (*52*).
- We could not find evidence of rabies infection in *Bos frontalis*, but as this is a semi-wild and endangered species (*53*) and other *Bos* species are susceptible, the disease may pose a conservation risk. This may also be the case for the endangered *Bos javanicus*.
- We did not find a specific report of rabies infection in *Vicugna vicugna*, although all South American camelids are noted to be susceptible and display clinical signs of infection (*54*).
- There are documented cases of rabies infecting *Rattus rattus* (ex. (*55*)), though it appears to be rare.
- Rabies in *Rattus norvegicus* was predicted by the affinity only model with the full dataset for domestic hosts.
- Pet guinea pigs *Cavia porcellus* have been infected with rabies after being bitten by a raccoon (*56*).
- There are reported cases of rabid *Oryctolagus cuniculus* in the United States (*22*).
- *Toxoplasma gondii* is known to infect *Bos grunniens* and cause severe economic losses (*57*).

### 2.9 Phylogeny only – Full Dataset – Wild Hosts

**Table SM 8:**
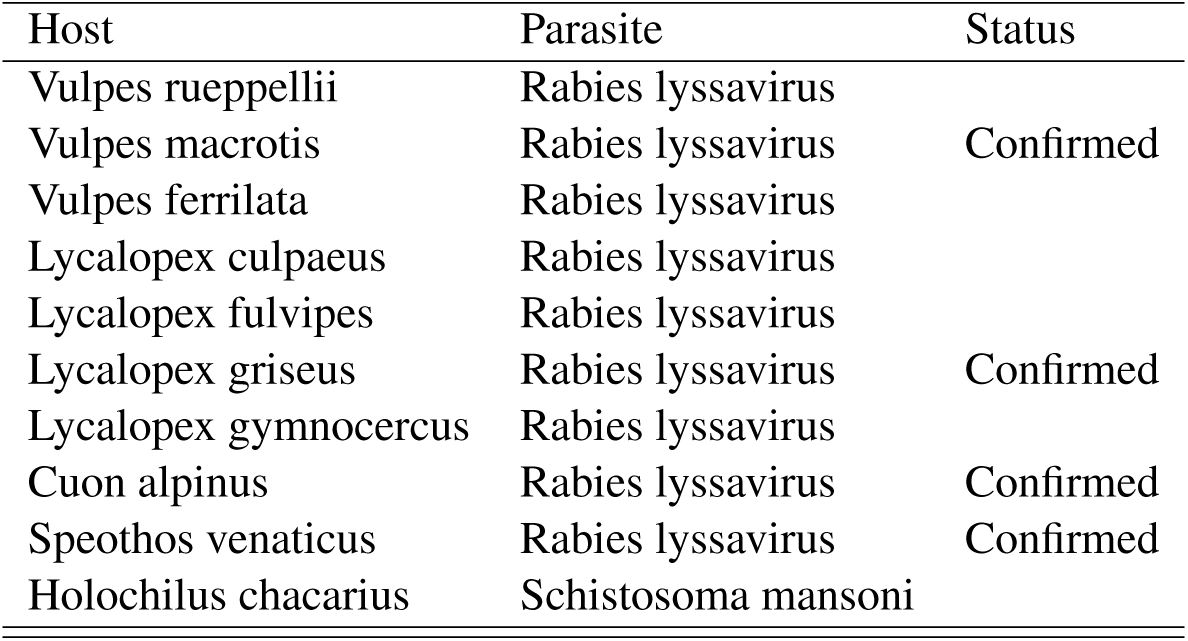
Top 10 undocumented links with highest probablity of interaction for wild host species: Phylogeny only model - full data set.

- These top predictions are the same as for the phylogeny only model on the full dataset (all wild host species in the top 10 links).

### 2.10 Combined model – Full Dataset

**Table SM 9:** Top 10 undocumented links with highest probablity of interaction (Combined model - full data set)

| Host | Parasite | Status |
| --- | --- | --- |
| Rattus norvegicus | Rabies lyssavirus | Confirmed |
| Cervus elaphus | Rabies lyssavirus | Confirmed |
| Homo sapiens | Taenia mustelae |  |
| Homo sapiens | Bluetongue virus |  |
| Ondatra zibethicus | Rabies lyssavirus | Confirmed |
| Capreolus capreolus | Rabies lyssavirus | Confirmed |
| Rattus rattus | Rabies lyssavirus | Confirmed |
| Sorex araneus | Rabies lyssavirus |  |
| Oryctolagus cuniculus | Rabies lyssavirus | Confirmed |
| Apodemus sylvaticus | Rabies lyssavirus | Confirmed |

- All of the top links were previously discussed except for Rabies infection of *Sorex araneus*, for which we could not find any published evidence.

### 2.11 Combined model – Full Dataset – Domestic Hosts

**Table SM 10:**
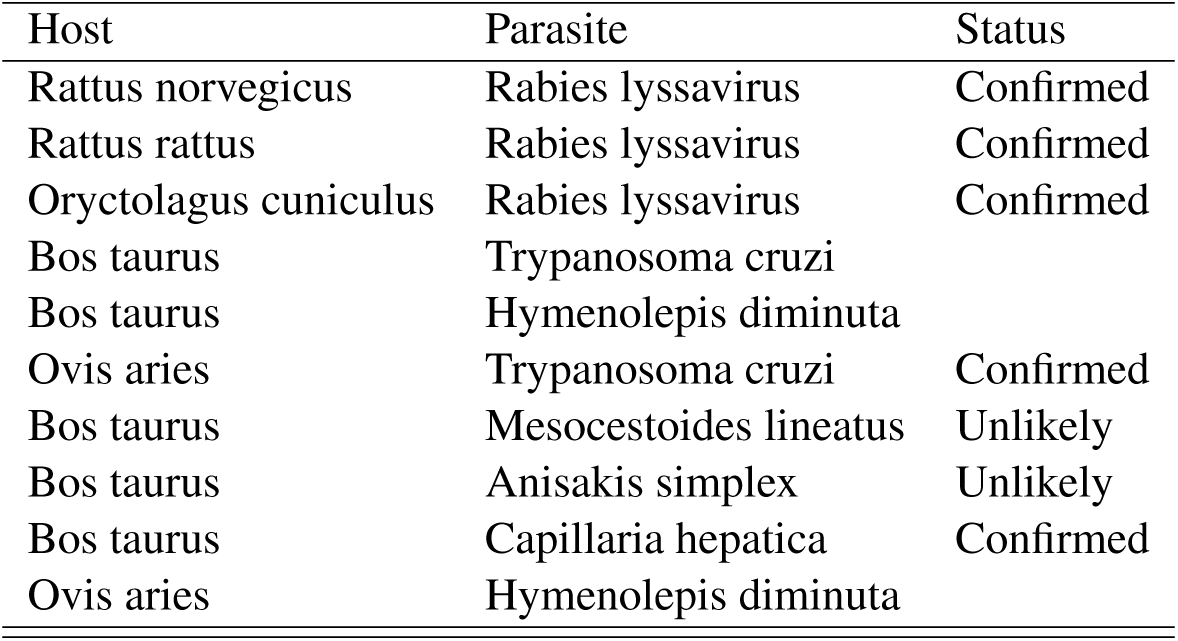
Top 10 undocumented links with highest probablity of interaction for domesticated species: Combined model - full data set.

- All of the top links were previously discussed except for *Hymenolepis diminuta* infection of sheep, for which we could not find any published evidence.

### 2.12 Combined model – Full Dataset – Wild Hosts

**Table SM 11:** Top 10 undocumented links with highest probablity of interaction for wild host species: Combined model - full data set.

| Host | Parasite | Status |
| --- | --- | --- |
| Cervus elaphus | Rabies lyssavirus | Confirmed |
| Ondatra zibethicus | Rabies lyssavirus | Confirmed |
| Capreolus capreolus | Rabies lyssavirus | Confirmed |
| Sorex araneus | Rabies lyssavirus |  |
| Apodemus sylvaticus | Rabies lyssavirus | Confirmed |
| Odocoileus virginianus | Rabies lyssavirus | Confirmed |
| Syncerus caffer | Rabies lyssavirus |  |
| Dama dama | Rabies lyssavirus | Confirmed |
| Aepyceros melampus | Rabies lyssavirus | Confirmed |
| Rupicapra rupicapra | Rabies lyssavirus | Confirmed |

- Most of these links were predicted by models discussed above except for rabies infection in *Aepyceros melampus*, which has been identified as suffering from spillover infections (*58*), and *Rupicapra rupicapra*, which has been documented in Europe (*36*).

### 2.13 Arthropods – Affinity only

**Table SM 12:** Top 10 undocumented links with highest probablity of interaction: Affinity only - arthropod subset.

| Host | Parasite | Status |
| --- | --- | --- |
| *Vulpes vulpes* | *Rhipicephalus evertsi* |  |
| *Vulpes vulpes* | *Rhipicephalus appendiculatus* |  |
| *Vulpes vulpes* | *Amblyomma hebraeum* |  |
| *Cervus elaphus* | *Rhipicephalus evertsi* |  |
| *Aepyceros melampus* | *Sarcoptes scabiei* | Confirmed |
| *Vulpes vulpes* | *Hyalomma truncatum* |  |
| *Cervus elaphus* | *Rhipicephalus appendiculatus* |  |
| *Sus scrofa* | *Rhipicephalus evertsi* |  |
| *Bos taurus* | *Sarcoptes scabiei* | Confirmed |
| *Odocoileus virginianus* | *Sarcoptes scabiei* |  |

- *Rhipicephalus evertsi* and *R. appendiculatus* are common ticks in East and Southern Africa (*59*) and are unlikely to interact with non-African hosts such as *Cervus elaphus* and *Vulpes vulpes* (though there are some populations of *Vulpes vulpes* in North Africa). Future iterations of our implemented link prediciton framework may benefit from the inclusion of information on geographic range overlap among host species, however this may reduce the ability to identify future host-parasite associations that may occur given range expansions or species translocations.
- *Amblyomma hebraeum*, the main vector of *Ehrlichia ruminantium* in southern Africa, prefer large hosts such as cattle and wild ruminants, although immature stages are found to feed on a wide range of hosts including scrub hares, guineafowl, and tortoises (*60*). Considering the species is restricted to southern Africa, it is unlikely to interact with Vulpes vulpes, although it’s wide host range during immaturity indicates that it may be suscpetible given the opportunity.
- (*61*) compiled a list of documented host species of sarcoptic mange (caused by *Sarcoptes scabiei*) which includes *Aepyceros melampus* and cattle (*Bos taurus*).
- *Hyalomma truncatum* is found across sub-Saharan Africa, where it commonly infests domestic and wild herbivores, and domestic dogs (*60*). Considering the species is restricted to Africa, it is unlikely to interact with Vulpes vulpes, although it’s tendency to infest domestic dogs indicates that it may be suscpetible given the opportunity.
- *Rhipicephalus appendiculatus* was found to be the most prevalent tick species on domestic pigs in the Busia District of Kenya (*62*), indicating that increased monitoring may also identify *R. evertsi* on domestic pigs.
- While multiple cervids have been reported with sarcoptic mange (*61*), we cannot find any report of infection in white-tailed deer *Odocoileus virginianus*, although there are numerous reports of infection with mange caused by *Demodex sp.*, including the host specific *Demodex odocoilei* (*63*), potentially indicating competition among *Sarcoptes* and *Demodex* species.

### 2.14 Arthropods – Phylogeny only

**Table SM 13:** Top 10 undocumented links with highest probablity of interaction: Phylogeny only model - Arthropod subset.

| Host | Parasite |  |
| --- | --- | --- |
| *Canis adustus* | *Sarcoptes scabiei* |  |
| *Lycalopex fulvipes* | *Sarcoptes scabiei* |  |
| *Lycalopex vetulus* | *Sarcoptes scabiei* |  |
| *Vulpes lagopus* | *Pulex irritans* |  |
| *Cerdocyon thous* | *Sarcoptes scabiei* | Confirmed |
| *Chrysocyon brachyurus* | *Sarcoptes scabiei* |  |
| *Cuon alpinus* | *Sarcoptes scabiei* | Confirmed |
| *Speothos venaticus* | *Sarcoptes scabiei* | Confirmed |
| *Vulpes macrotis* | *Dermacentor variabilis* | Confirmed |
| *Ovis ammon* | *Sarcoptes scabiei* |  |

- There does not appear to be a published record of sarcoptic mange in *Canis adustus*, however in areas with sympatric jackal species *C. adustus* usually display ecological segregation through preferring denser vegetation (*64*). This may indicate that while *C. adustus* may be susceptible to sarcoptic mange, differences in the ecologies of this species relative to other canids may limit transmission making overt infections difficult to document.
- *Lycalopex fulvipes* is endangered (*44*), meaning that its small population sizes and restricted geographic range may reduce exposure to *S. scabiei*, however as sarcoptic mange is implicated in the declines of other wild canids, it should be targeted in disease monitoring programs for this species.
- While *Lycalopex vetulus* is not endangered, it displays some adaptability to anthropogenic disturbance (*65*), which may expose it to sarcoptic mange through contact with domestic dogs. In addition, *Lycalopex vetulus* is sympatric with the crab eating fox (*Cerdocyon thous*) – a documented host of *S. scabiei* (*61*). The IUCN reports a gap in conservation actions for *L. vetulus* regarding the role of disease in population regulation, and their status as reservoirs of scabies, canine distemper, leishmaniasis, and rabies (*65*).
- While the Arctic fox (*Vulpes lagopus*) is considered the most important terrestrial game species in the Arctic (*66*), we were unable to find documented infection by the “human flea” (*Pulex irritans*). *P. irritans* is thought to be unable to persist in Arctic envrionments due to the temperature thresholds necessary for breeding (*67*), though this may change in the future with continued Arctic warming.
- (*61*) compiled a list of documented host species of sarcoptic mange (caused by *Sarcoptes scabiei*) which includes *Cerdocyon thous*.
- *Chrysocyon branchyurus* has one report of clinical signs suggestive of sarcoptic mange-like infestation (*68*).
- The endangered Dhole (*Cuon alpinus*) has been documented as suffering from mange as early as 1937 (*47*) and appear to be especially susceptible to disease outbreaks due to their large group sizes and amicable behaviour within packs.
- *S. scabei* was identified in *Speothos venaticus* (*69*), and identified as potentially contributing to the loss of individuals from a group in Mato Grosso, Brazil (*70*).
- *Vulpes macrotis* was documented as infested with *Dermacentor variabilis* in Southwest Texas in 1950 (*71*).
- We cannot find a record of mange in *Ovis ammon*, though outbreaks of sarcoptic mange have been documented in ibex and blue sheep in the Taxkorgan Reserve, China, in which *O. ammon* are also present, although this population has received little study (*72*).

### 2.15 Arthropods – Combined model

**Table SM 14:** Top 10 undocumented links with highest probablity of interaction: Combined model - arthropod subset.

| Host | Parasite | Status |
| --- | --- | --- |
| Cerdocyon thous | Sarcoptes scabiei | Confirmed |
| Cervus elaphus | Rhipicephalus evertsi |  |
| Vulpes velox | Sarcoptes scabiei |  |
| Aepyceros melampus | Sarcoptes scabiei | Confirmed |
| Vulpes macrotis | Sarcoptes scabiei | Confirmed |
| Chrysocyon brachyurus | Sarcoptes scabiei |  |
| Odocoileus virginianus | Rhipicephalus evertsi |  |
| Odocoileus virginianus | Sarcoptes scabiei |  |
| Lycalopex vetulus | Sarcoptes scabiei |  |
| Bos taurus | Sarcoptes scabiei | Confirmed |

- We were unable to find any published evidence of *Sarcoptes scabiei* infesting *Vulpes velox*. However, sarcoptic mange is known to infest several species of canids, is prevalent in coyotes (*Canis latrans*) within the range of *Vulpes velox* (*73*). Criffield2009 surveyed for *S. scabiei* on *V. velox*, but no clinical signes were observed, and suggest that as the grey fox (*Urocyon cineroargenteus*) is somewhat resistant to mange in laboratory tests, *V. velox* may be similarly resistant, though there have been no experimental infestations. Finally, *V. velox* is debated to be conspecific with *Vulpes macrotis* (*74*), which has been documented with sarcoptic mange (*75*).
- *Sarcoptes scabiei* has been documented in the endangered San Jaoaquin kit fox *Vulpes macrotis mutica*, and is considered a significant threat to it’s conservation (*75*).
- *Rhipicephalus evertsi* is a common tick in East and Southern Africa (*59*) and is unlikely to interact with non-African hosts such as *Odocoileus virginianus*. Future iterations of our link prediction method may benefit from the inclusion of information on geographic range overlap among host species, however this may reduce the ability to identify future host-parasite associations that may occur given range expansions or species translocations.
- The remaining links are discussed above.

### 2.16 Bacteria – Affinity only

**Table SM 15:** Top 10 undocumented links with highest probablity of interaction: Affinity only model - bacteria subset.

| Host | Parasite | Status |
| --- | --- | --- |
| Homo sapiens | *Bartonella grahamii* | Confirmed |
| Homo sapiens | *Anaplasma marginale* |  |
| Homo sapiens | *Anaplasma bovis* |  |
| Homo sapiens | *Mycoplasma mycoides* |  |
| Homo sapiens | *Mycoplasma haemofelis* |  |
| Homo sapiens | *Chlamydophila pecorum* |  |
| Homo sapiens | *Lawsonia intracellularis* |  |
| Homo sapiens | *Mycoplasma conjunctivae* |  |
| Homo sapiens | *Chlamydia psittaci* | Confirmed |
| Homo sapiens | *Histophilus somni* |  |

- *Bartonella grahamii* is a pathogen of rodents worldwide, but was first identified as causing an infection in an immunocompromised human in 2013 (*76*).
- *Anaplasma bovis*, causal agent of bovine anaplasmosis, is not currently considered zoonotic (*77*), but *Anaplasma phagocytophilum* the causative agent of human anaplasmosis is placed is as sister taxa to *A. bovis* in a recent phylogeny (*78*). Similarly, *A. marginale*, the causative agent of anaplasmosis in cattle, is also not considered zoonotic, but it reaches high prevalence in cattle and humans are likely exposed to the tick vector (*77*).
- *Mycoplasma mycoides* is not typically thought to infect humans, but there is one report of disease and positive serology in a farm worker exposed to multiple calves infected with *M. mycoides subsp. mycoides LC* (*79*).
- There has been one documented case of infection in an immunocompromised human with a *Mycoplasma haemofelis*-like bacteria (*80*) and the authors note that disease-causing latent mycoplasma infections in immunocompromised and non-immunocompromised patients are an emerging issue.
- The zoonotic potential of *Chlamydophila pecorum* is not known, although it is associated with abortions in small ruminants (*81*).
- *Lawsonia intracellularis* was recently recognised as the cause of an emerging intestinal disease in horses (Equine proliferative enteropathy), but is currently not considered to be zoonotic (*82*).
- *Mycoplasma conjunctivae* causes a highly contagious ocular infection of sheep, goats, and wild Caprinae, and is possibly zoonotic as it has been associated with eye inflammation in young children (*83*).
- *Chlamydophila psittaci* (formerly known as *Chlamydia psittaci*) is a known zoonotic disease from birds (*81*).
- *Histophilus somni* is a pathogen of bovine and ovine hosts, but has not been documented to infect humans (*84*).

### 2.17 Bacteria – Phylogeny only model

**Table SM 16:**
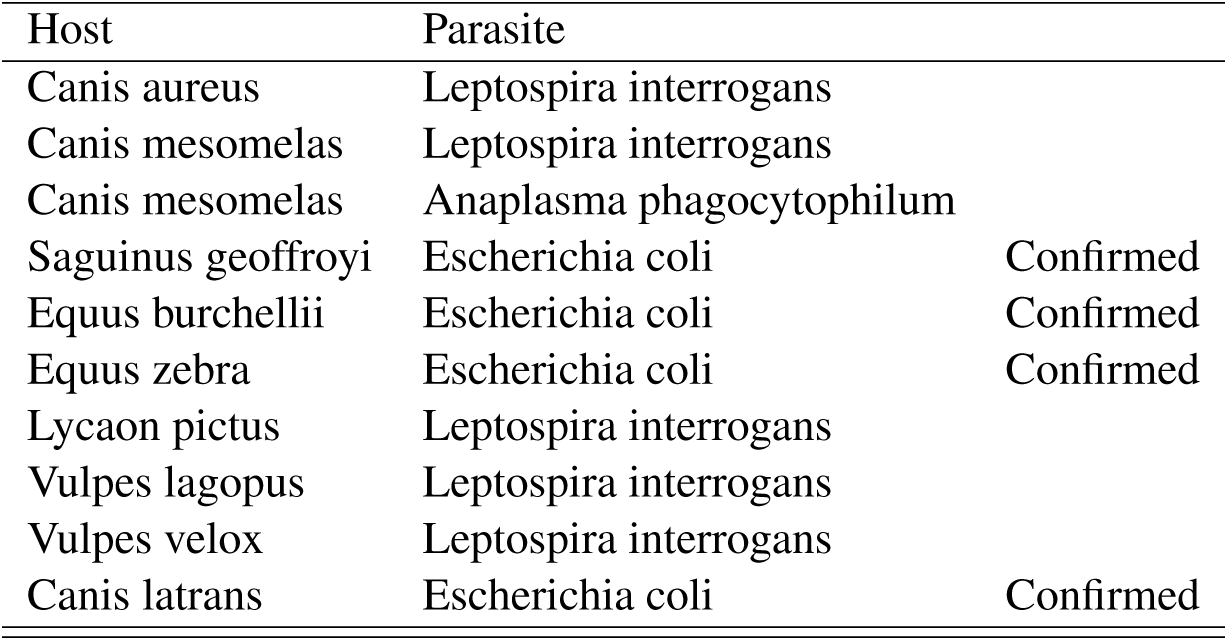
Top 10 undocumented links with highest probablity of interaction: Phylogeny only - bacteria subset.

- *Leptospira sp.* are commonly regarded as infecting a wide range of mammals (*85*). (*86*) identified *Leptospira interrogans* serovar *canicola* in the urine of jackals in Israel though did not identify the particular species. However, *Canis aureus* is the only jackal species present in the country (*87*) providing some support for this host-parasite association, though the findings of (*86*) should be verified.
- We did not find any documentation of leptospirosis in *Canis mesomelas* or *Lycaon pictus* but considering it is found in multiple species in Africa including domestic dogs (*88*), wild canids are likely to be exposed.
- (*89*) sequenced DNA from blood samples of *Canis mesomelas* in South Africa and identified 16S rDNA sequences very similar to *Anaplasma phagocytophilum*. This study also identified other *Anaplasma* species indicating the potential for 16S rDNA sequencing to gather evidence of predicted host-parasite and discover previously unknown pathogens.
- *E. coli* is a ubiquitous commensal microbe of vertebrates (*90*) and pathogenicitiy is linked to particular strains, indicating that our approach may be expanded by identifying the host ranges of particular subspecies or virulent strains of common commensal bacteria. *Canis latrans* has been identified as harbouring atypical enteropathogenic *E. coli* and may serve as a reservoir in agricultural areas near the United States-Mexico border (*91*). Wild Equus burchellii have been found to harbour antibiotic resistant *E. coli* in South Africa (*92*) and Tanzania (*93*), and captive *Equus zebra hartmannae* have been found with antibiotic resistant *E. coli* (*94*). Similarly, antibiotic resistant *E. coli* have been found in captive *Saguinus geoffroyi* (*95*).
- We did not find any evidence of *Leptospira* infections in *Vulpes lagopus*. We did find one report of positive serology for *Leptospira interrogans* in *Vulpes velox macrotis* (*96*), though there is debate as to whether this subspecies is actually its own species *Vulpes macrotis* (*1*).

### 2.18 Bacteria – Combined model

**Table SM 17:** Top 10 undocumented links with highest probablity of interaction: Combined model - bacteria subset.

| Host | Parasite | Status |
| --- | --- | --- |
| Ovis canadensis | Escherichia coli |  |
| Bison bison | Escherichia coli | Confirmed |
| Ovis aries | Mycobacterium bovis | Confirmed |
| Homo sapiens | Bartonella grahamii | Confirmed |
| Ovis canadensis | Leptospira interrogans |  |
| Zalophus californianus | Anaplasma phagocytophilum | Unlikely |
| Ovis canadensis | Anaplasma phagocytophilum |  |
| Phoca vitulina | Leptospira interrogans |  |
| Homo sapiens | Anaplasma marginale |  |
| Bison bison | Anaplasma phagocytophilum |  |

- *E. coli* is ubiquitous commensal microbe of vertebrates (*90*) and *Ovis canadensis* has been surveyed for pathogenic *E. coli* in Washington, USA, though no individuals tested positive (*97*). *Bison bison* have been highlighted as a potentially important reservoir of pathogenic *E. coli O157:H7* for human infection (*98*).
- While there is some debate whether sheep are relatively immune or highly susceptible to infection by *Mycobacterium bovis*, spillover infections have been documented to occur when animals are exposed to contaminated pasture (*99*).
- *Ovis canadensis* has been identified as exposed to *Leptospira interrogans* through serological surveys (*100*).
- While tick-borne *Anaplasma phagocytophilum* is found to persist in a large range of terrestrial mammalian hosts (*101*), we find no evidence of infection in *Zalophus californianus* or any other marine mammals.
- We identified one survey of *Anaplasma sp.* in Ovis canadensis in Montana which conducted testing for *Anaplasma phagocytophilum*, though no individuals were positive (*102*). (*102*) speculate that this lack of infection despite the potential for exposure may be due to the exclusion of Anaplasma genotypes such as A. ovis, which is found to infect *Ovis canadensis*.
- Regular exposure of *Phoca vitulina* to *Leptospira interrogans* has been reported (*103*), though low antibody titers were interpreted as exposure rather than infection.
- We did not find evidence of *Anaplasma phagocytophilum* infection in American bison (*Bison bison*), though *A. phagocytophilum* is known to infect European bison (*Bison bonasus*) in Poland (*101*).
- The remaining links are discussed above.

### 2.19 Fungi – Affinity only model

**Table SM 18:**
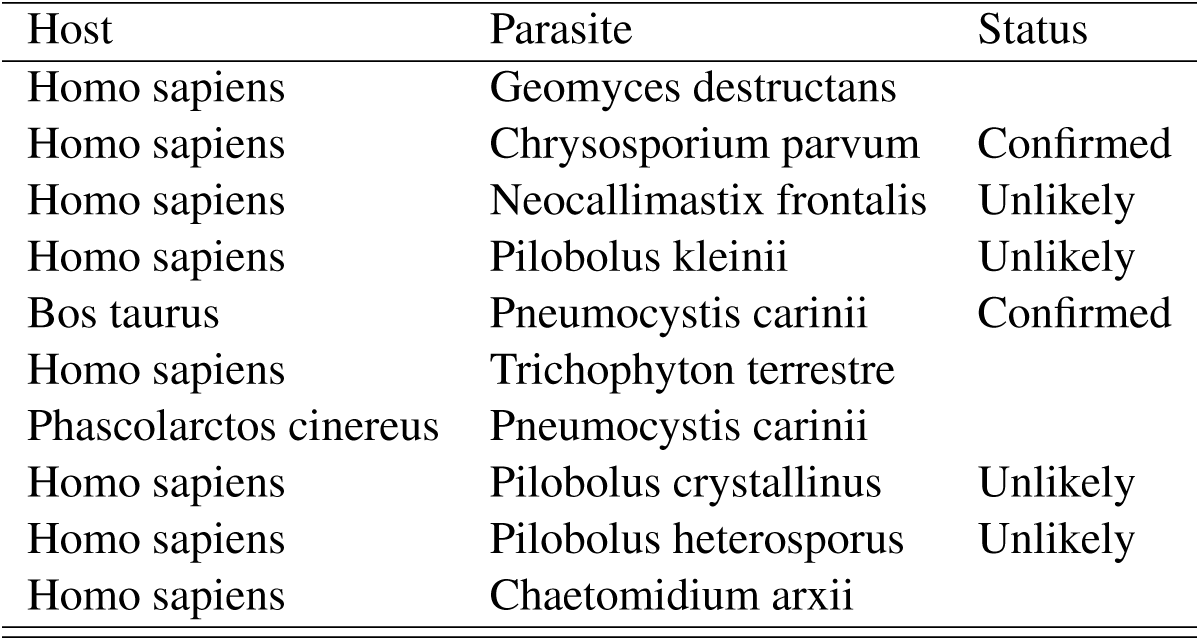
Top 10 undocumented links with highest probablity of interaction: Affinity only - fungi subset.

- *Geomyces destructans* is the cause of white nose syndrome in multiple bat species (*104*), but is not considered to be zoonotic.
- *Chrysosporium parvum* and related species are soil fungi that cause pulmonary infections in rodents, fossorial mammals, their predators, and occasionally humans, though the taxonomy of these pathogens is muddied in the literature (*105*).
- *Neocallimastix frontalis* appears to be a commensal fungi of bovid rumens (*106*) and we cannot find documentation of zoonotic infection.
- *Pilobolus sp.* play a role in the decomposition of herbivore dung and although they are nonpathogenic to herbivores, they can facilitate the spread of attached parasitic lungworms because of their projectile dispersal system (*107*). We could find no evidence of human infections.
- *Pneumocystis carinii* belongs to a genus that normally reside in the pulmonary parenchyma of a wide range of mammals (*108*). It is capable of causing life threatening pneumonia in immuno-compromised hosts and is documented as causing infections in cattle (*109*), though we found no evidence of infection in *Phascolarctos cinereus*.
- *Trichophpyton terrestre* is part of a large species complex with some variants documented to cause human infection (*110*).
- We cannot find any documentation of *Chaetomidium arxii* infection in humans, although this genus is well known for its opportunistic animal and human pathogens (*111*).

### 2.20 Fungi – Phylogeny only

**Table SM 19:** Top 10 undocumented links with highest probablity of interaction: Phylogeny only - fungi subset.

| Host | Parasite | Status |
| --- | --- | --- |
| Macaca sylvanus | Pneumocystis carinii |  |
| Felis silvestris | Pneumocystis carinii |  |
| Mustela lutreola | Pneumocystis carinii |  |
| Ateles paniscus | Pneumocystis carinii |  |
| Pan troglodytes | Pneumocystis carinii |  |
| Gorilla beringei | Pneumocystis carinii |  |
| Mustela erminea | Pneumocystis carinii |  |
| Ovis canadensis | Pneumocystis carinii |  |
| Nyctereutes procyonoides | Pneumocystis carinii |  |
| Vulpes vulpes | Pneumocystis carinii | Confirmed |

- *Pneumocystis carinii* belongs to a genus that normally reside in the pulmonary parenchyma of a wide range of mammals (*108*). It is capable of causing life threatening pneumonia in immuno-compromised hosts is documented as causing infections cattle and pigs (*112*), *Vulpes vulpes* (*113*), and species in the genera *Macaca* and *Mustela* (*113*).

### 2.21 Fungi – Combined model

**Table SM 20:**
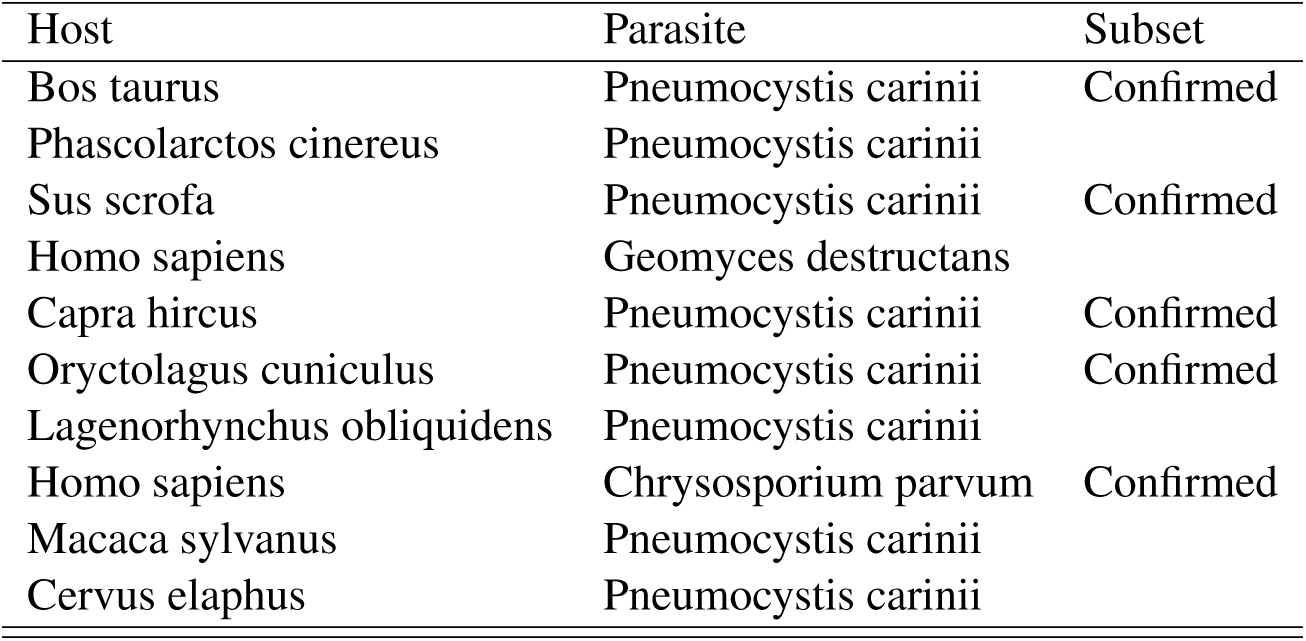
Top 10 undocumented links with highest probablity of interaction: Combined model - fungi subset.

- *Pneumocystis carinii* has been documented to infect pigs (*112, 108*), goats (*108*), and *Oryctolagus cuniculus* (*113*). While it has been documented in other cervids (*113*), we find no evidence of infection in *Cervus elaphus*.
- The remaning top links are discussed above.

### 2.22 Helminths – Affinity only model

**Table SM 21:**
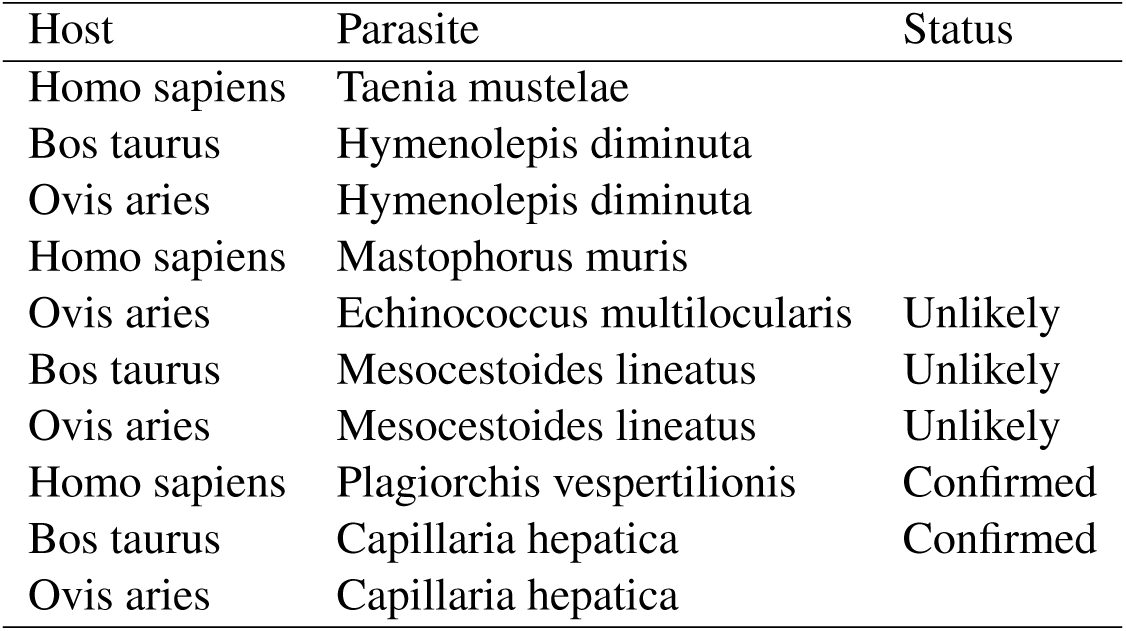
Top 10 undocumented links with highest probablity of interaction: Affinity only - helminth subset.

- Although the distribution, ecology, and epidemiology of *Echinococcus multilocularis* in North America is still largely unknown, it does not appear to infect sheep or any other ungulates as it is maintained in a carnivore-rodent prey cycle (*114*).
- *Mesocestoides lineatus* has a three-stage lifecycle with two intermediate hosts and a large range of carnivorous mammals as definitive hosts (*24*). Human infections of the tapeworm *Mesocestoides lineatus* are rare but can occur through the consumption of chickens, snails, snakes, or frogs (*25*) and therefore it is unlikely that sheep will consume the intermediate life stages of this parasite.
- *Capillaria hepatica* (syn. *Calodium hepaticum*), is a globally distributed zoonotic parasite which uses rodents as main hosts, and while it is known to cause infection in over 180 mammalian species, a recent review of hosts did not include domesticated sheep (*26*). However, a recent study of *Capillaria* in Brazil identified two cases which were possibly caused by *C. hepatica* (*115*).
- The remaning links are discussed above.

### 2.23 Helminths – Phylogeny only

**Table SM 22:** Top 10 undocumented links with highest probablity of interaction: Phylogeny only - helminth subset.

| Host | Parasite | Status |
| --- | --- | --- |
| Holochilus chacarius | Schistosoma mansoni |  |
| Onychogalea unguifera | Echinococcus granulosus |  |
| Canis adustus | Echinococcus granulosus |  |
| Gazella leptoceros | Nematodirus spathiger | Confirmed |
| Kobus vardonii | Cotylophoron cotylophorum |  |
| Onychogalea unguifera | Rugopharynx australis |  |
| Lycalopex vetulus | Echinococcus granulosus | Unlikely |
| Canis mesomelas | Trichinella spiralis | Confirmed |
| Gazella leptoceros | Trichostrongylus vitrinus |  |
| Onychogalea fraenata | Progamotaenia festiva |  |

- *Schistosoma mansoni* infection in *Holochilus chacarius* was predicted by the phylogeny only model in the full dataset (discussed above).
- *Echinococcus granulosus* has not been reported to infect northern nail-tail wallabies (*Onychogalea unguifera*), other wallaby species including endangered bridled nail-tailed wallaby (*Onychogaela fraenata*) are involved in the transmission the parasite in Australia (*116*). *Onychogaela unguifera* may also be involved in *Echinococcosus* transmission, but its parasites may not be as well studied compared to the bridled nail-tail wallaby due to its stable conservation status.
- Similarly, *Rugopharynx australis* is known to infect multiple wallaby species, however the diversity of *Rugopharynx* and their susceptible hosts is still being discovered (*117*), suggesting that *Onychogalea ungifera* may be a promising target for future study.
- While there does not appear to be evidence of *Echinococcosus granulosus* infection in *Canis adustus*, other *Canis* species in Africa are known hosts (*118*), indicating that this should be a target for future surveillance.
- A recent molecular survey of gastrointestinal parasites of wild ruminants in Tunisia identified *Nematodirus spathiger* in engandered *Gazella leptoceros* that were genetically identical to those found in other domestic and wild ruminants (*119*). This is an example of a successful exploratory study aimed at describing the diversity of parasites in threatened species.
- We did not find much information on the parasites of the near threatened *Kobus vardonii*, however it is known to inhabit floodplains and grasslands near permanent water in south-central Africa (*120*) where is likely to be exposed to *Cotylophoron cotylophoron*, a “rumen fluke” which emerge from snail intermediate hosts and encyst on vegetation, later being ingested by ruminant definitive hosts in East Africa (*121*).
- *Echinococcus granulosus* is usually maintained by a domestic cycle of dogs eating raw livestock offal (*118*), and while its vertebrate-eating congener *Lycalopex gymnocercus* has been documented to host the parasite (*122*), *Lycalopex vetulus* is unlikely to become infected with *E. granulosus* as it has a largely insectivorous diet (*123*).
- *Canis mesomelas* has been reported with infection of *Trichinella spiralis* in the Kruger National Park, South Africa (*124*).
- *Gazella leptoceros* is also predicted to be susceptible to *Trichostrongylus vitrinus*. Although (*119*) did not identify this parasite in their study, *T. vitrinus* has been documented in lambs in Tunisia (*125*), indicating potential range overlap with *G. leptoceros*.
- *Progamotaenia festiva* is known to infect multiple *Onychogalea* species (*126*), but we could not find evidence of infection in the endangered *Onychogalea fraenata*.
- The remaining links are discussed above.

### 2.24 Helminths – Combined model

**Table SM 23:** Top 10 undocumented links with highest probablity of interaction: Combined model - helminth subset.

| Host | Parasite | Subset |
| --- | --- | --- |
| Homo sapiens | Taenia mustelae |  |
| Bos taurus | Hymenolepis diminuta |  |
| Ovis aries | Hymenolepis diminuta |  |
| Ovis aries | Echinococcus multilocularis | Unlikely |
| Capra hircus | Hymenolepis diminuta |  |
| Bos taurus | Mesocostoides lineatus | Unlikely |
| Ovis aries | Mesocostoides lineatus | Unlikely |
| Bos taurus | Anisakis simplex | Unlikely |
| Ovis aries | Anisakis simplex | Unlikely |
| Sorex araneus | Echinococcus granulosus |  |

- The natural hosts of *Hymenolepis diminuta* are rats (*23*) and we find no evidence of infection in goats.
- *Anisakis simplex* uses cetaceans as final hosts, with marine invertebrates and fish as intermediate hosts (*30*). Whales are infected through ingestion, indicating that while sheep may be susceptible, though they would need sufficient exposure to marine based feed.
- We could not find evidence of *Echonococcus granulosus* infection in Sorex araneus, however this species and other *Sorex sp.* are known hosts of *Echinococcus multilocularis* (*127*).
- The other top links are discussed above.

### 2.25 Protozoa – Affinity only model

**Table SM 24:**
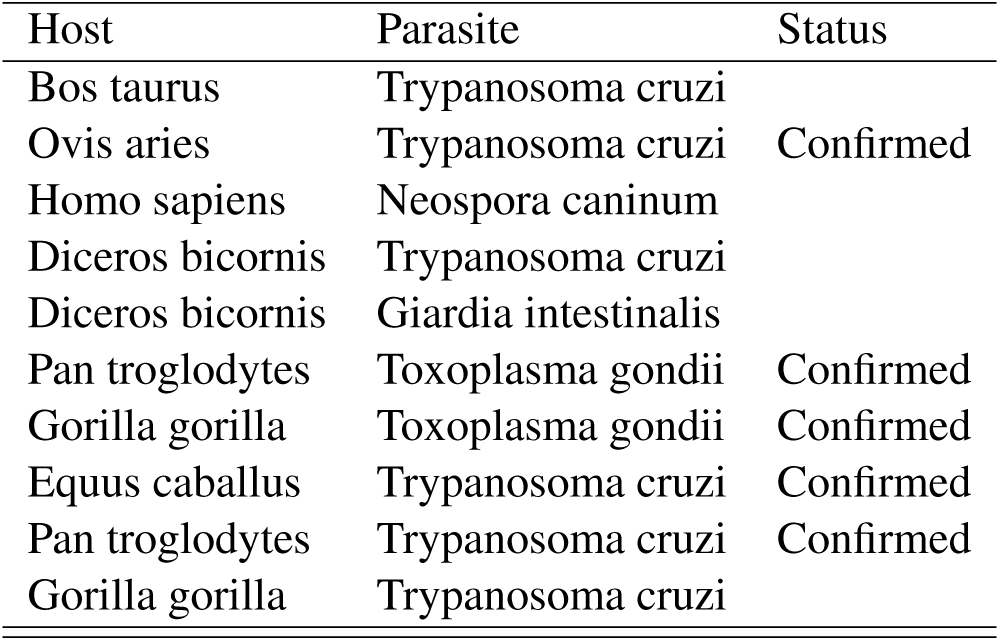
Top 10 undocumented links with highest probablity of interaction: Affinity only - protozoa subset.

- As *T. cruzi* is currently restricted to the Americas (*20*), it is unlikely to infect black rhinos (*Diceros bicornis*) or gorillas (*Gorilla gorilla*) in natural conditions, unless facilitated by human activities.
- *Giardia* has been identified in a captive bred *Diceros bicornis* calf (*128*) in San Diego, indicating the potential for grey literature from zoo and captive breeding facilities to inform potential host-parasite interactions.
- *Toxoplasma gondii* has been documented to infect chimpanzees (*Pan troglodytes*), and interestingly appears to mirror the infection-induced behaviour in rodents and humans, with infected chimpanzees attracted to the urine of leopards, their only natural predator (*129*).
- Recent finding of a *Gorilla gorilla* individual seropositive for *T. gondii* at a primate center in Gabon (*130*).
- *T. cruzi* was recently identified in *Equus caballus*, marking the first evidence of infection in equids (*131*).
- Although *T. cruzi* naturally occurs in the Americas, and thus natural infection of chimpanzees (*Pan troglodytes*) is unlikely, a fatal infection was documented in a captive individual in Texas (*132*).

### 2.26 Protozoa – Phylogeny only

**Table SM 25:** Top 10 undocumented links with highest probablity of interaction: Phylogeny only - protozoa subset.

| Host | Parasite | Subset |
| --- | --- | --- |
| Canis aureus | Toxoplasma gondii |  |
| Saguinus oedipus | Trypanosoma cruzi | Confirmed |
| Cuon alpinus | Toxoplasma gondii |  |
| Lycaon pictus | Toxoplasma gondii | Confirmed |
| Saimiri oerstedii | Trypanosoma cruzi | Confirmed |
| Mazama gouazoubira | Toxoplasma gondii | Confirmed |
| Nyctereutes procyonoides | Toxoplasma gondii | Confirmed |
| Saguinus niger | Trypanosoma cruzi | Confirmed |
| Capricornis swinhoei | Toxoplasma gondii |  |
| Panthera tigris | Toxoplasma gondii | Confirmed |

- *Canis aureus* with antibodies against *T. gondii* have been identified in captive animals in the United Arab Emirates (*133*).
- Two recent reviews of parasites in non-human primates find no documented infection of the critically endangered Cotton-top tamarin (*Saguinus oedipus*) by *Trypanosoma cruzi*, although multiple *Saguinus sp.* have been documented with infections (*134, 135*). However, a 1982 study of Colombian monkeys and marmosets identified *S. oedipus* as a host for *T. cruzi* for the first time (*136*). This highlights the potential conservation importance of this parasite for *S. oedipus* and the need for periodic disease surveys of critically endangered species.
- Similarly, *Saimiri oerstedii* is not listed by these reviews as a host of *T. cruzi*, although a 1972 study identifies *S. oerstedii* as a reservoir for the parasite in Panama (*137*). While this report should be followed up with contemporary diagnostic methods, this reiterates the difficulty of exhaustively searching the literature for interaction data and the utility of link prediction methods to for directing these efforts.
- *Toxoplasmoa gondii* infection in *Cuon alpinus* has rarely been investigated, except for one captive individual which tested negative in serological testing (*138*).
- High prevalence of antibodies against *Toxoplasma gondii* was found in wild dogs (*Lycaon pictus*) in the Kruger National Park, South Africa, and was documented as causing a fatal infection in one pup (*139*), indicating that this parasite has the potential to influence the population dynamics of this endangered canid.
- *T. gondii* has been identified in *Mazama gouanzoubira* from French Guiana (*140*) and *Nyctereutes procyonoides* (*141*) in China via genetic sequencing.
- *Trypanosoma cruzi* is documented to infect *Saguinus niger* (*135*).
- We did not find any reports of *T. gondii* infection in Taiwan serow *Capricornis swinhoei*, although direct evidence of infection has been found in Japanese serow (*142*) and *T. gondii* has been found to infect multiple animals in Taiwan (*143*).
- The Siberian tiger *Panthera tigris altaica* acts as a definitive host for *T. gondii* and is observed to naturally shed oocysts (*144*).

### 2.27 Protozoa – Combined model

**Table SM 26:**
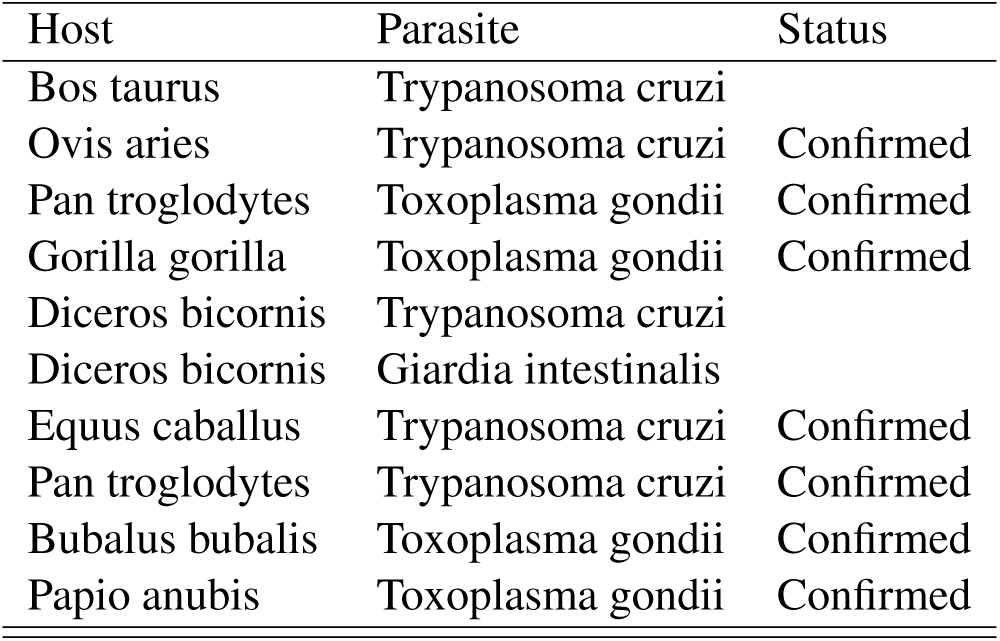
Top 10 undocumented links with highest probablity of interaction: Combined model - protozoa subset.

- The first eight links were predicted by models discussed above.
- *Bubalus bubalis* is a well established host of *Toxoplasma gondii*, though they are considered resistant to clinical toxoplasmosis (*145*).
- Natural infections of *Toxoplasma gondii* in *Papio anubis* were recently documented via genetic sequencing (*146*).

### 2.28 Viruses – Affinity only

**Table SM 27:** Top 10 undocumented links with highest probablity of interaction: Affinity only - virus subset.

| Host | Parasite | Subset |
| --- | --- | --- |
| Homo sapiens | Bluetongue virus |  |
| Homo sapiens | Bovine viral diarrhea virus 1 |  |
| Homo sapiens | Carnivore protoparvovirus 1 |  |
| Homo sapiens | Simian immunodeficiency virus | Confirmed |
| Pan troglodytes | Rabies lyssavirus | Confirmed |
| Macaca mulatta | Rabies lyssavirus | Confirmed |
| Homo sapiens | Bovine alphaherpesvirus 1 |  |
| Homo sapiens | Canine mastadenovirus a |  |
| Cervus elaphus | Rabies lyssavirus | Confirmed |
| Homo sapiens | Ovine herpesvirus 2 |  |

- Simian immunodificiency virus strains from wild primates have previously shifted to infect humans and are responsible for the AIDS pandemic (from HIV-1) (*147*).
- Rabies positive *Macaca mulatta* have recently been reported in India (*148*).
- Bovine alphaherpesvirus 1, the main casual agent of infectious bovine rhinotracheitis, is largely restricted to cattle and not currently considered to infect humans (*149*).
- *Canine mastadenovirus a*, formerly *Canine adenovirus 1* is known to infect dogs and circulate in wild carnivores (*15*), though we could not find evidence of human infection.
- *Ovine herpesvirus 2*, the casual agent of sheep associated malignant catarrhal fever, is asymptomatic in its natural hosts and cause severe disease in susceptible animals that are dead end hosts, however to date there is no evidence of infection in humans (*150*).
- The remaining links are discussed above.

### 2.29 Viruses – Phylogeny only

**Table SM 28:** Top 10 undocumented links with highest probablity of interaction: PHylogeny only - virus subset.

| Host | Parasite | Status |
| --- | --- | --- |
| *Vulpes macrotis* | Rabies lyssavirus | Confirmed |
| *Lycalopex culpaeus* | Rabies lyssavirus |  |
| *Lycalopex griseus* | Rabies lyssavirus | Confirmed |
| *Lycalopex gymnocercus* | Rabies lyssavirus |  |
| *Myotis nattereri* | Rabies lyssavirus | Confirmed |
| *Myotis blythii* | Rabies lyssavirus | Confirmed |
| *Myotis myotis* | Rabies lyssavirus | Confirmed |
| *Myotis macrodactylus* | Rabies lyssavirus |  |
| *Myotis mystacinus* | Rabies lyssavirus |  |
| *Myotis dasycneme* | Rabies lyssavirus | Confirmed |

- The first four links were predicted by previous models and discussed above.
- Rabies has been isolated from a single *Myotis nattereri* individual in France (*151*).
- In 2017, an individual *Myotis blythii* from Croatia tested positive for antibodies against rabies (*152*).
- Rabies has been detected in *Myotis myotis* in a few European countries (*153*).
- We did not find evidence of rabies infection in *Myotis macrodactylus* or *Myotis mystacinus*.
- Identification of rabies positive *Myotis dasycneme* in the Netherlands (*154*).

### 2.30 Viruses – Combined model

**Table SM 29:**
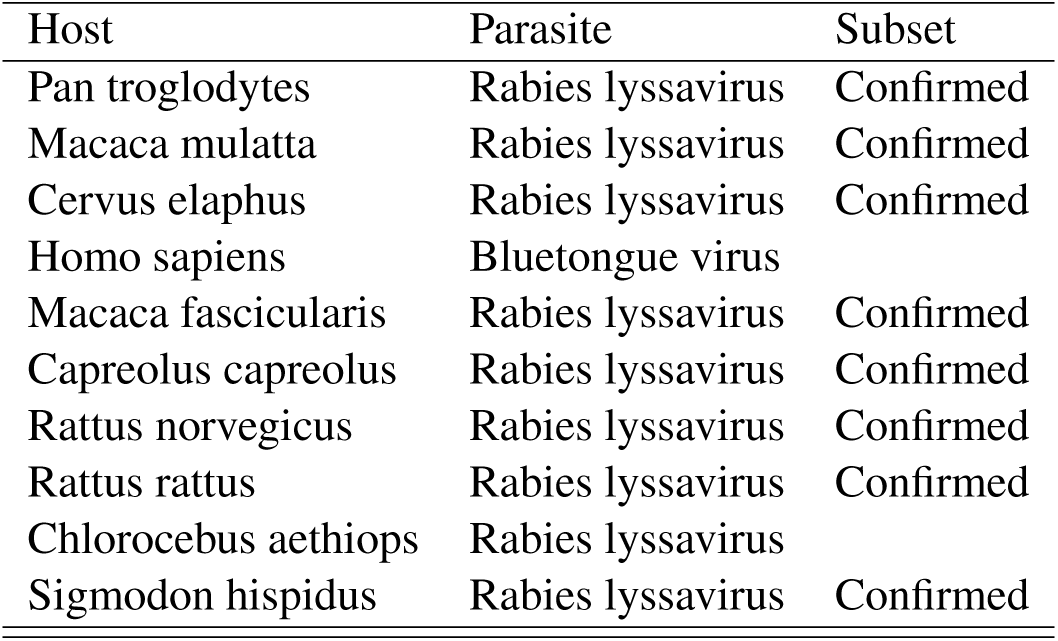
Top 10 undocumented links with highest probablity of interaction: Combined model - virus subset

| Host | Parasite | Subset |
| --- | --- | --- |
| Pan troglodytes | Rabies lyssavirus | Confirmed |
| Macaca mulatta | Rabies lyssavirus | Confirmed |
| Cervus elaphus | Rabies lyssavirus | Confirmed |
| Homo sapiens | Bluetongue virus |  |
| Macaca fascicularis | Rabies lyssavirus | Confirmed |
| Capreolus capreolus | Rabies lyssavirus | Confirmed |
| Rattus norvegicus | Rabies lyssavirus | Confirmed |
| Rattus rattus | Rabies lyssavirus | Confirmed |
| Chlorocebus aethiops | Rabies lyssavirus |  |
| Sigmodon hispidus | Rabies lyssavirus | Confirmed |

- *Macaca fascicularis* is known to be susceptible to infection by *Rabies lyssavirus* and regularly used for development of rabies vaccines (*155*).
- We did not find evidence of rabies infection in *Chlorcebus aethiops*, however several cercopithecine monkeys are known to be susceptible to infection (*35*).
- *Sigmodon hispidus* has successfully been experimentally infected with rabies (*156*).
- The remaining links are discussed above.

## Notes

http://doi.org/10.6084/m9.figshare.8969882

http://github.com/melmasri/HP-prediction

### Competing Interest Statement

The authors have declared no competing interest.

### Summary of Updates

Small changes to the text throughout, additional analyses, one revised figure.

